# Plasticity of fibroblast transcriptional response to physical and biochemical cues revealed by dynamic network analysis

**DOI:** 10.1101/2020.12.13.422572

**Authors:** Pilhwa Lee, Joseph Decker, Lonnie Shea, Daniel A. Beard

## Abstract

Data on human skin fibroblast transcriptional responses to external cues were used to reconstruct dynamic gene regulatory networks. The goal of the reconstruction was to determine dynamic network interactions (quantitative predictive relationships of mutual regulatory influences of and on transcription factor expression) from time course data on 56 transcript expression levels obtained following different external cues. The inherently under-determined nature of this problem was addressed in part by excluding putative regulatory motifs that did not appear to be functional in multiple independent experiments from different independent external perturbations. Data were obtained from a previously published experiment in which the 56 transcripts were assayed by bioluminescence in live cells cultured on substrates of varying levels of stiffness and exposed to different levels of arginylglycylaspartic acid (RGD) peptide. The inferred dynamical networks were validated via comparison of predictions to *a priori* known interactions from gene databases. We discovered that exposures to different substrate stiffnesses and to RGD stimulate responses that are mediated through *GATA4, SMAD3/4, ETS-1*, and *STAT5* and other genes, which can initiate hypertrophic, fibrotic, and inflammatory responses. The developed dynamical system identification method for discovering new mechanotransduction pathways is applicable to the identification of gene regulatory networks in numerous emerging applications where time-series data on multiple state variables and from multiple external perturbations are available.

## Introduction

Fibroblasts, which are widely present in many tissue types, contribute to the regulation of the extracellular environment by secreting constituent proteins of the extracellular matrix. By influencing the mechanical properties of the microenvironment, fibroblasts play essential roles in numerous physiological, developmental, and pathophysiological disease processes. Specific developmental and disease processes in which fibroblast functions are known to make important contributions are cell lineage specification (*1-10*), epithelial-mesenchymal interaction (*11-13*), and epithelial-mesenchymal transition (*14-18*) as well as carcinogenesis and metastasis (*19-31*) and immunogenic plasticity in inflammatory diseases (*32-35*). In each of these settings, the ways in which fibroblasts respond to particular mechanical and biochemical stimuli determine how they operate. For example, with certain cardiac-specific factors as biochemical stimuli, fibroblasts may be reprogrammed to cardiac progenitors with multi-potency to cardiomyocytes, smooth muscles, and endothelial cells (*8*). Following an epithelial injury in the lung, fibroblasts respond by secretion of pro-fibrotic factors and can be transdifferentiated to myofibroblasts accompanied by ECM stiffening and fibrosis (*33*). Associated with triple-negative breast cancer, fibroblasts may also be transdifferentiated to myofibroblastic cancer-associated fibroblasts (CAF’s). One sub-type of CAF induces regulatory T cells to inhibit T effector proliferation in heterogeneity (*31*) and promotes cancer invasion (*25*).

To understand the mechanisms underlying the plasticity of fibroblast responses to different stimuli, Shea et al. used the state-of-the-art live-cell imaging technique, TRAnscriptional Factor CEll ARray (TRACER) (*36*), to obtain time-series data on multiple transcripts from human skin fibroblasts following perturbations in extracellular matrix (ECM) stiffness and arginylglycylaspartic acid (RGD) peptide levels. RGD peptide is ubiquitously responsible for cell adhesion to the ECM. Thus, by perturbing RGD peptide levels and ECM stiffness, fibroblasts are exposed to a range of mechanical stimuli designed to yield a rich, dynamic response in cellular transcriptional states. To probe the system’s interactions in the complex response observed by Shea et al., we used a dynamic network modeling approach to investigate the transcriptional response by attempting to reconstruct network-level regulatory pathways with multiple time-point data from the live-cell experiments of Peñalver Bernabé et al. (*36*).

Most approaches in RNA-level transcriptomic investigation are designed to analyze static data from a single time point. Approaches to network inference from time-series data include Bayesian inference, linear regression, and mutual information techniques (*37-41*). Recent advances in the single-cell analysis have generated dynamic models by incorporating a non-physical “pseudo time” to represent changes following a perturbation (*42-45*). Here, we apply a method designed to infer a “physical-time” differential equation-based model for transcription factor expression and regulation from time-series data on multiple transcripts (*46, 47*). We deconvolve gene-to-gene connectivity treating the transcriptomic ensemble as a kinetic dynamical system on randomized networks and projecting to the empirical time-series data from Shea et al. (*36*). Theoretically, this problem is a non-polynomial (NP) hard, that is, generally not possible to solve exactly. We have approximately handled this difficulty by decomposing a network into subnetworks with one-to-many relationships utilizing GPU-ODE based parallel reconstruction of subnetworks (*46, 47*). Those sub-networks are compared to kinetic data by a hybrid approach of global and gradient-based optimizations. By scoring gene-to-gene interaction and deriving *p*-values from multiple conditions, we identify commonly inferred directly associated genes and differentially inferred pathways from external differential cues. The selection of random networks for testing is unbiased in bootstrapping. i.e., there is no incorporation of *a priori* confirmed gene-to-gene interactions. The result is a set of likely dynamic network interactions that are informed solely by the time-course data and not by prior knowledge.

Our analysis focuses on approximately 60 transcriptional factors mainly involved in proliferation, differentiation, apoptosis, chemical and hormonal receptors. Based on the gene regulatory network (GRN) architecture inferred from the mechano-transduction data, we identify several putative differential mechanisms by which master regulatory genes *GATA4, SMAD3/4, ETS1*, and *STAT5* relay information from upstream signals to hypertrophic, fibrotic, or inflammatory phenotypes diversely. The influences of ECM stiffness and cell-ECM adhesion on the regulation of these genes are cooperative in some cases and counter-cooperative in others. Validation of these results is based on *a priori* known interactions from published databases. The validated results reveal the promise of this approach for larger-scale kinetic network reconstruction.

## Methods

### Transcriptional activity measurement, TRACER

The expression of transcriptional factor activities of human skin fibroblast was measured at five time points (3, 6, 9, 12, 27 hours) by the bioluminescent optical technique TRACER (*TR*anscriptional *A*ctivity *CE*ll a*R*rays). Six different external conditions were prescribed by three different RGD levels at one PEG hydrogel stiffness as well as three different PEG hydrogel stiffnesses with one RGD level in combination. For RGD variant conditions, RGD concentrations are prescribed as 0.7, 2.0, and 3.3 mM with hydrogel stiffness 1.5 kPa. For hydrogel stiffness variant conditions, PEG-hydrogel stiffnesses are prescribed as 0.15 kPa, 1.5 kPa, and 6.5 kPa with RGD concentration of 2.0 mM. For full experimental details, see Peñalver Bernabé et al. (*36*).

### Gene regulatory network reconstruction

The process of reconstructing fibroblast gene regulatory network takes five steps: (1.) Clustering, (2.) Identification of sub-networks, (3.) Development of consensus model from sub-network models, (4.) Global network construction, and (5.) Network validation by *a priori* known databases (Figure 1).

**Figure 1:**
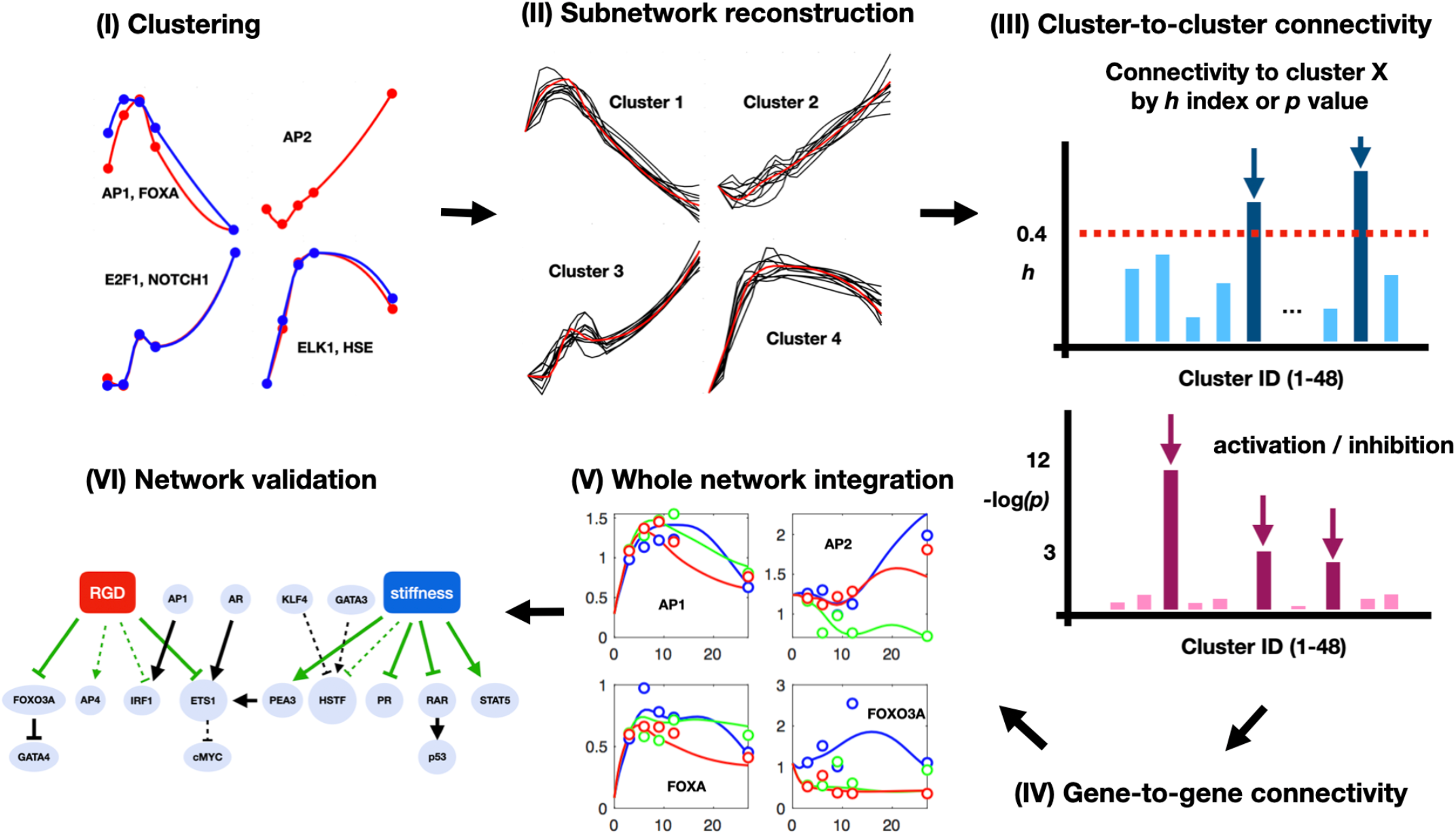
Workflow of gene network reconstruction. **(I) Clustering**. Four clusters are shown as samples in RGD-high and stiffness-medium condition. **(II) Subnetwork reconstruction**. Acceptable subnetworks are reconstructed for the four sampled clusters in (I). **(III) Cluster-to-cluster connectivity**. Significance of connectivity to each specific cluster X is determined by *h*-indices or *p*-values. **(IV) Gene-to-gene connectivity** Cluster-to-cluster connectivity is decompressed to gene-to-gene connectivity. **(V) Whole network integration** Time course profiles of each gene expression in 6 conditions are fitted in global optimization with fixed *h*-index based gene-to-gene connectivity to identify cue-to-gene interactions. Fitting to four sample gene profiles is shown with three differential RGD levels. **(VI) Network validation** *P*-value based significant cue-to-gene interactions (green) and gene-to-gene (black) interactions are classified as valid (solid) or predicted (dotted)) based on published literature (Table 1).

#### 1. Clustering

For each of the six independent experiments representing different external conditions, the time courses of the 56 measured transcripts were compared to one another to identify clusters with similar behavior. For a given experiment, we denote the measured data for transcript *i* at time point *t*_*k*_ as *x*_*i*_(*t*_*k*_). The first step in clustering is to normalize the data

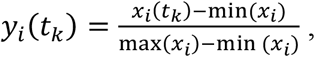

where min(*x*_*i*_) and max(*x*_*i*_) are the minimum and maximum values of *x*_*i*_ over the time course.

The clustering of transcript time courses for a given experimental condition is based on several independent metrics of similarity. The first metric, *D*_1_(*i, j*), is the temporal correlation of transcriptional factors *i* and *j* defined:

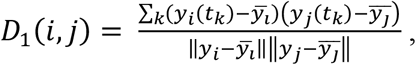

where 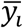 is the mean value of *y*_*i*_(*t*_*k*_) over a given time course. This calculation is a normalized inner-product of *y*_*i*_(*t*_*k*_) and *y*_*j*_(*t*_*k*_) with their mean values as baselines.

The second metric is a measurement of similarity in the local slopes of *y*_*i*_(*t*_*k*_) and *y*_*j*_(*t*_*k*_):

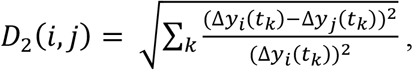

where Δ*y*_*i*_(*t*_*k*_) = *y*_*i*_(*t*_*k*+1_) − *y*_*i*_(*t*_*k*_).

The third metric of similarity is based on the numbers of convex and concave critical points estimated to exist in the time courses for *y*_*i*_(*t*_*k*_) and *y*_*j*_(*t*_*k*_). Convex critical points are identified at times *t*_*k*_ when *y*_*i*_(*t*_*k*−1_) > *y*_*i*_(*t*_*k*_) and *y*_*i*_(*t*_*k*_) < *y*_*i*_(*t*_*k*+1_). Similarly, concave critical are identified at times *t*_*k*_ when *y*_*i*_(*t*_*k*−1_) < *y*_*i*_(*t*_*k*_) and *y*_*i*_(*t*_*k*_) > *y*_*i*_(*t*_*k*+1_).

Given these three metrics, two time profiles from transcription factors *i* and *j* are clustered together when: (1.) the indicator of temporal correlation *D*_1_(*i, j*) > *α*; (2.) the indicator of differences in temporal gradients *D*_2_(*i, j*) < *β*; and (3.) the number of critical points in *y*_*i*_(*t*_*k*_) and *y*_*j*_(*t*_*k*_) are the same. For the calculations presented here, the parameters *α* and *β* are set to 0.9 and 0.45, respectively.

The original 56 gene time courses were clustered into 47 clusters for the RGD-high and stiffness-medium experiment, 45 clusters for the RGD-middle and stiffness-medium experiment, 46 clusters for the RGD-low and stiffness-medium, 37 clusters for the RGD-middle and stiffness-hard experiment, 44 clusters for the RGD-middle and stiffness-medium experiment, and 37 clusters for the RGD-middle and stiffness-soft experiment. Clusters for the six experiments are illustrated in Supplementary Figures S1-S3.

#### 2. Identification of sub-networks

Expression of transcripts is assumed to be governed by the following general model (*46*):

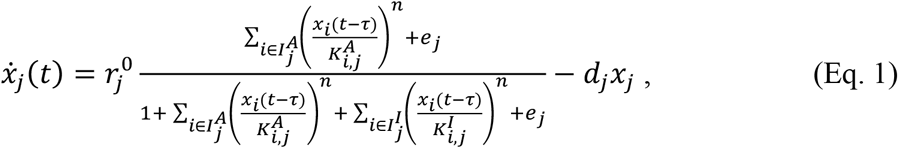

where the terms in the expression are defined as follows:

*x*_*j*_(*t*): mRNA expression of gene *j* at time *t*

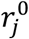: basal transcription rate of gene *j*

*d*_*j*_: mRNA degradation rate of gene *j*

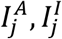: a set of indices of activators and inhibitors on gene *j*

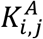: dissociation constant for activation of gene *j* by gene *i*

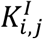: dissociation constant for inhibition of gene *j* by gene *i*

*e*_*j*_: external signals on gene *j*

*τ*: time delay between transcription and translation

*n*: cooperative Hill-coefficient

The model assumes competitive and cooperative binding of inhibitory and activating factors. The time delay constant *τ* and the Hill-coefficient *n* are prescribed as 1 hour and 2, respectively. The other parameters, activation/inhibition indices 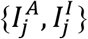 and kinetics parameters 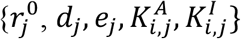, are estimated by fitting with the experimental data.

The details of the model identification process are described in detail in (*46*). In brief, the model of Equation (1) is applied to each cluster, rather than at the individual gene level. For a given cluster for a given experiment, the governing equation is treated as a one-dimensional (one state variable) ordinary differential equation for a given variable *x*_*j*_(*t*) by using shape-preserving piecewise cubic interpolations of the measured data for other variables *x*_*i*_(*t*) on the right-hand side of Equation (1). Each of these one-dimensional problems is highly underdetermined with multiple solutions that can equivalently match the observed data for a given variable *x*_*j*_(*t*). The algorithm works by performing a massively parallel search for model structures (defined by the activation/inhibition indices 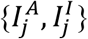) and parameter values 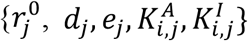 associated with each putative structure. Using a GPU architecture, the algorithm returns >100 putative models at once for each cluster for each experimental condition.

Figure 2 shows time-course data associated with 16 example clusters (panel A, red lines) obtained from the RGD-high and stiffness-medium condition. The total number of clusters attained for this experiment was 47. For each cluster profile, 100 regulatory sub-networks that are able to match the cluster time-course within the error threshold are obtained from randomized bootstrapping and GPU-ODE integration. Panel B of Figure 2 shows the best five and worst five model fits from the total ensemble of sub-network model fits for each of these clusters.

**Figure 2.**
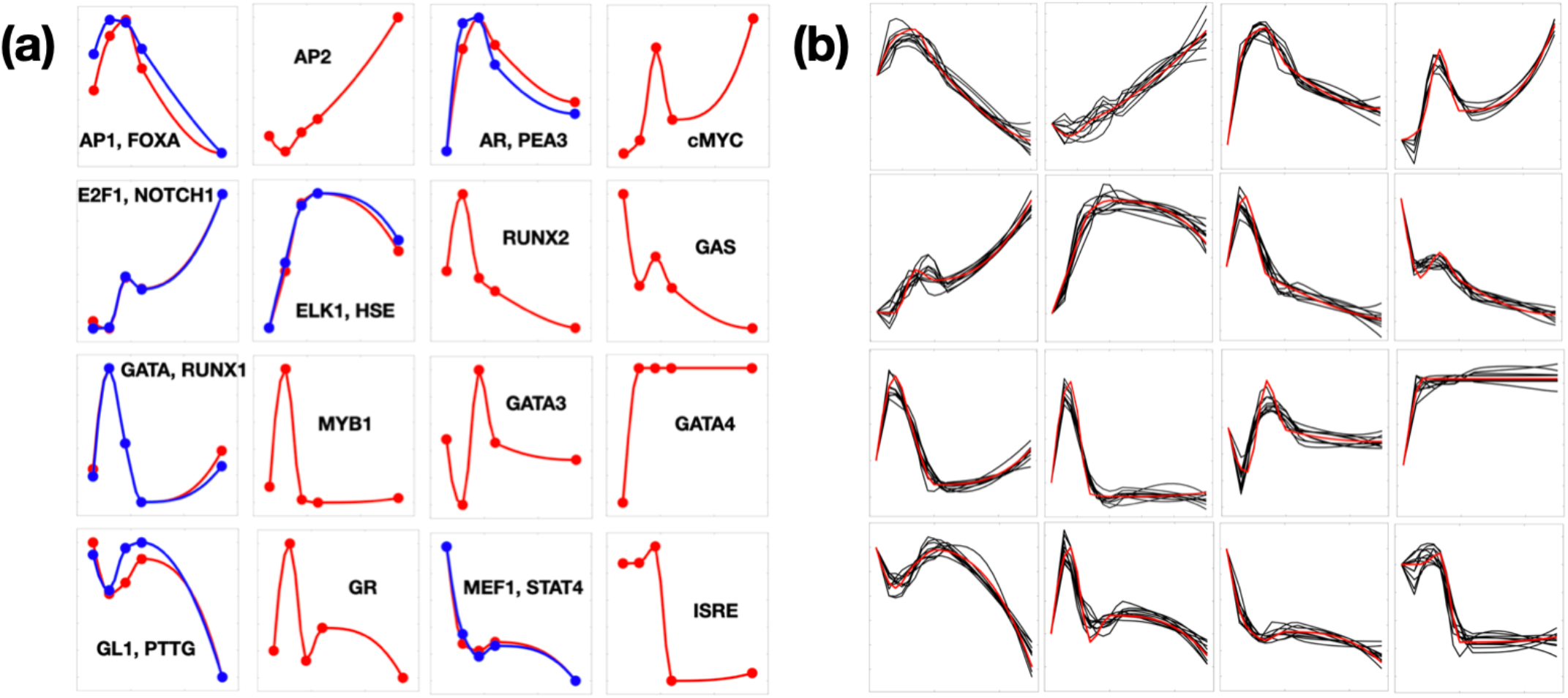
Gene profile clustering and subnetwork reconstruction by randomized connectivity. (a) Clustering. The blue curved profiles are clustered to each red profile as a unique feature based on correlation and occurrences of convexity. Each dataset from 6 conditions of external cues has different clustering profiles. (b) Reconstruction of subnetworks. From each cluster profile, a subnetwork is reconstructed with randomized bootstrapping by GPU-ODE integration and 100 times fitting within the error threshold. For the time course profiles averaged among the clustered genes from experiments (red), best five and worst five fitted profiles generated by the kinetic model on random networks (black) are shown together.

#### 3. Development of consensus model from sub-network models

Step 2 (identification of sub-networks) yields sets of putative activation and inhibition interactions between genes in the clusters. Each cluster-cluster interaction represents *n* × *m* potential gene-gene interactions, where *n* and *m* are the numbers of genes in each cluster. To judge the likelihood of a putative gene-gene interaction, we first enumerate the *n* × *m* potential interactions for each cluster-cluster interaction in the set of sub-networks identified for all clusters from all experiments.

We assess the likelihood of a given identified gene-gene interaction representing a biologically real interaction based on the frequency with which that interaction appears in subnetworks identified from the six independent experiments. Our underlying assumption is that the greater the frequency with which a given interaction appears, the more likely that interaction represents real biological functions. For each putative interaction, we estimate the equivalent of an “h-index” citation index, *h*. Here, the factor *h* is defined as the fraction of experiments for which a given interaction appears with a frequency equal to *h*. For example, if for a given interaction *h* = 0.25, then this interaction appears with a frequency of 25% or greater in 25% or more subnetworks identified for the independent experiments.

In each condition, the h-index of statistical fidelity of activation from gene *i* to gene *j* is defined as (the occurrence of accepted random networks with activation from gene *i* to gene *j*) / (total occurrence of accepted random networks). The factor of *h* for inhibition from gene *i* to gene *j* is similarly calculated. The overall h-index is calculated by fitting the sorted collection of six individual h-indices from six conditions to 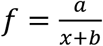, and taking the shortest distance from the origin to the fitted reciprocal curve (Figure 3a). Panels b, c, and d of Figure 3 illustrate the overall network connectivity obtained with different thresholds for the value of the h-index. For the lowest value of the index illustrated (panel b, *h* = 0.24) there are a total of 533 activation and inhibition interaction in the network of 56 genes. For an intermediate value of the index illustrated (panel c, *h* = 0.26) there are a total of 248 activation and inhibition interaction in the network of 56 genes. For the highest value of the index illustrated (panel d, *h* = 0.28) there are a total of 103 activation and inhibition interaction in the network of 56 genes.

**Figure 3.**
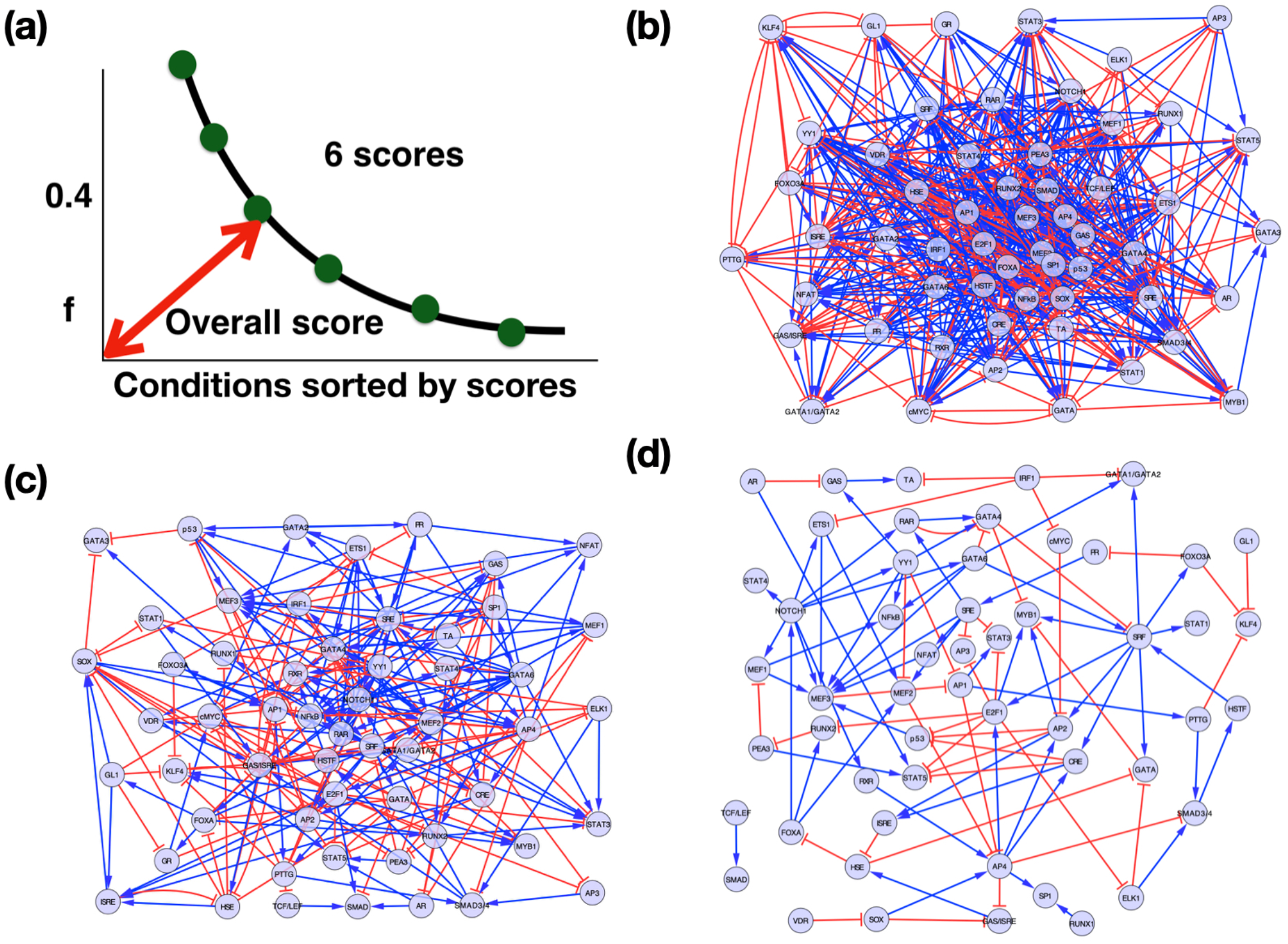
Global gene network integration by scoring from multiple cues. (a) Gene-to-gene fidelity is estimated from gene-to-gene binding frequency-based score combining six scores as one. (b) For those gene-to-gene edges that scored more than the threshold of overall *h*=0.24, the gene-to-gene interactions of the activation (blue) and inhibition (red) are prescribed for the global network. (c) The gene regulatory network with the overall *h*-index threshold=0.26. (d) The gene regulatory network with the overall *h*-index threshold=0.28.

For each external condition, we also define the significance of gene-to-gene interactions based on estimated *p*-values. The *p*-values are calculated from statistical distribution of gene-to-gene binding occurrence events (Figure 4a and 4c) following the steps:

**Figure 4.**
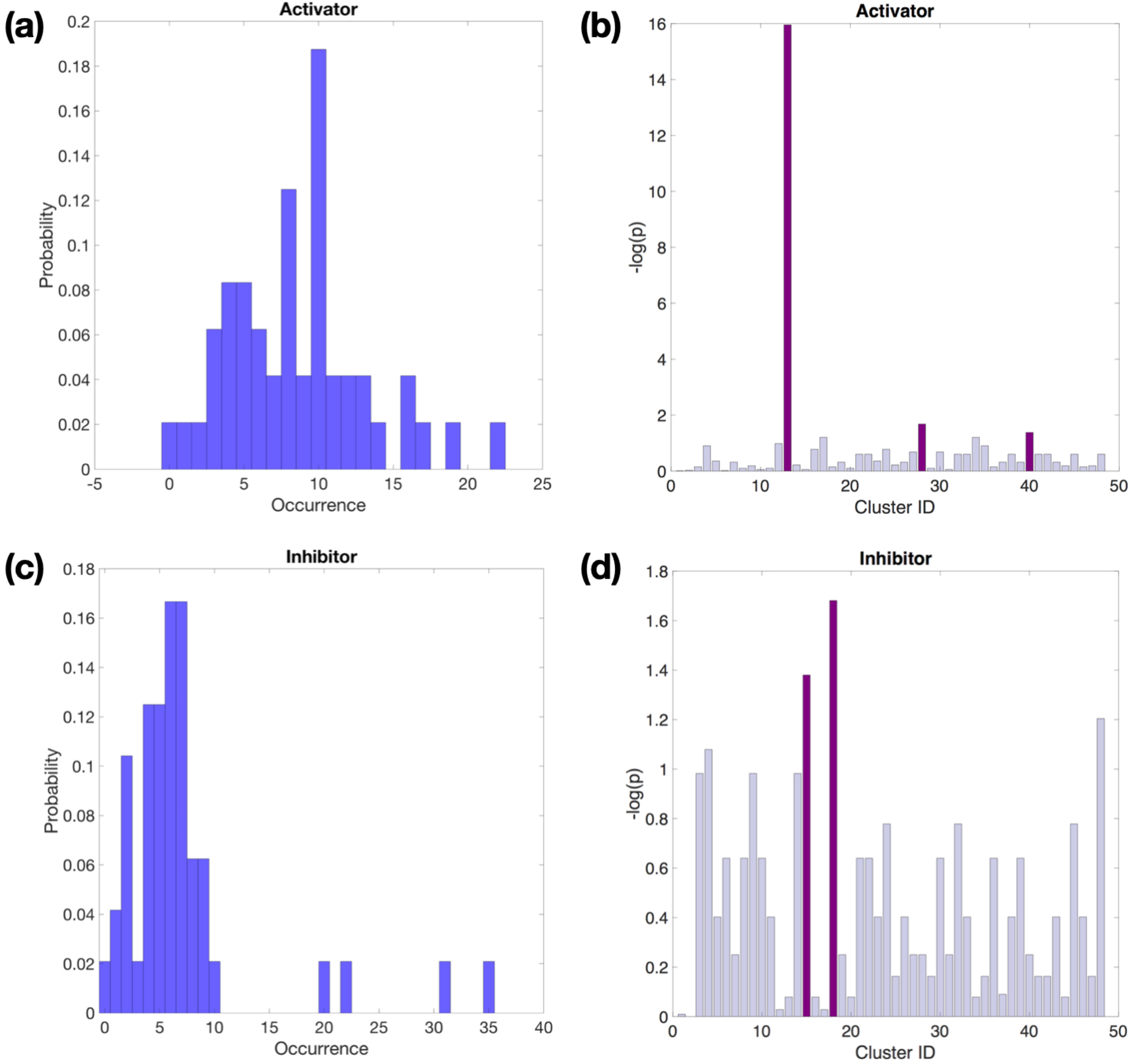
Reconstruction of p-value based gene regulatory subnetworks. The subnetwork reconstruction process generates unbiased random gene binding events of activation/inhibition for each cluster in each condition. The binding occurrences are collected as distribution (a and c), and the clusters are identified as significant for *p* < 0.05 (b and d) and colored as violet. The insignificant clusters are colored as light violet.

1. Based on cluster-to-cluster interactions, the occurrences of activation and inhibition interactions are enumerated, from which a frequency distribution of occurrences of interaction is obtained. For example, Figure 4a shows about 8 clusters function with 10 occurrences of activation on a target cluster in 100 acceptable subnetworks with total 45 clusters. Similarly, Figure 4c shows that about 1 cluster functions with 35 occurrences of inhibition on the target cluster in 100 acceptable subnetworks with total 45 clusters. There is a mapping *M*(*i, j*) from each cluster *i* to its occurrence of interaction with the specific cluster *j*.
2. From these frequency distributions of occurrences, we derive cumulative probability density distributions for activation and inhibition interactions, respectively.
3. For each interaction from cluster *i* to cluster *j* (activation or inhibition), the occurrence of interaction *M*(*i, j*) points to the corresponding cumulative probability density “*p*_cummulation_”.
4. The cluster-to-cluster interactions are identified as significant for *p* = 1-*p*_cummulation_< 0.05. Activators and inhibitors identified as significant are highlighted as violet in Figure 4 b and d. Insignificant clusters are colored as light violet.
5. The *p* values of *n* × *m* gene-gene interactions are obtained from the corresponding cluster-cluster interaction.

#### 4. Global network construction

For a given assumed cutoff value for the frequency parameter *h*, we obtain a set of internal network interactions defining the global network for the 56 transcription factors studied here. Figure 3 shows examples of network with 533 edges (*h* = 0.24), 248 edges (*h* = 0.26), and 103 edges (*h* = 0.28). The lower the cutoff *h* value, the more interaction edges accepted into the network.

Our goal is to simulate and explain whole-network behavior in terms of Equation (1) where each transcript is treated as a dependent variable, rather than in terms of one-variable sub-networks. To do this, we need to obtain global estimates of the activation and inhibition dissociation constants associated with each activation and inhibition interaction in the network, as well as the external signal factors for each gene for each of the six experimental conditions.

To simulate the global network in an integrated manner the external cues to 56 genes are modeled by a linear combination of saturating effects from two independent cues from RGD peptide and PEG-hydrogel stiffness:

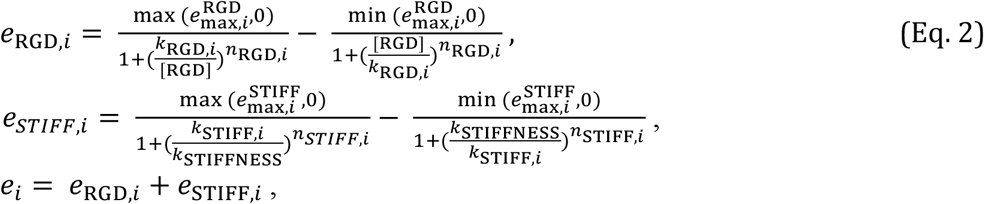

where 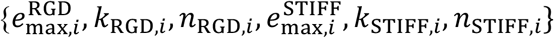 are estimated by fitting whole-network function to the time-course data for all genes from all external cue conditions. For a given gene *i* the external cues *e*_RGD,*i*_ and *e*_STIFF,*i*_ can represent either activation or inhibition depending on the sign of the parameters 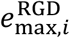 and 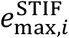.

Because the experiments have no data for *t* = 0, we obtain a long-time steady state for the isolated system ([RGD] = 0, *k*_STIFFNESS_ = 0) to use at the initial condition for each simulation. To estimate global kinetic parameters, the 56 gene profiles from 6 conditions are fit using a combination of stochastic simulated annealing, gradient-based optimization, and the globally convergent method of moving asymptotes (GCMMA) (*48*). Using this hybrid approach, we obtain a number of independent realization of network parameter sets associated with eight relatively equivalent fits to the data (Figure S7-S14). Figures 5 show examples time-course fits for 16 gene transcripts for differential RGD levels (panel a) and differential hydrogel stiffness levels (panel b) experiments.

**Figure 5.**
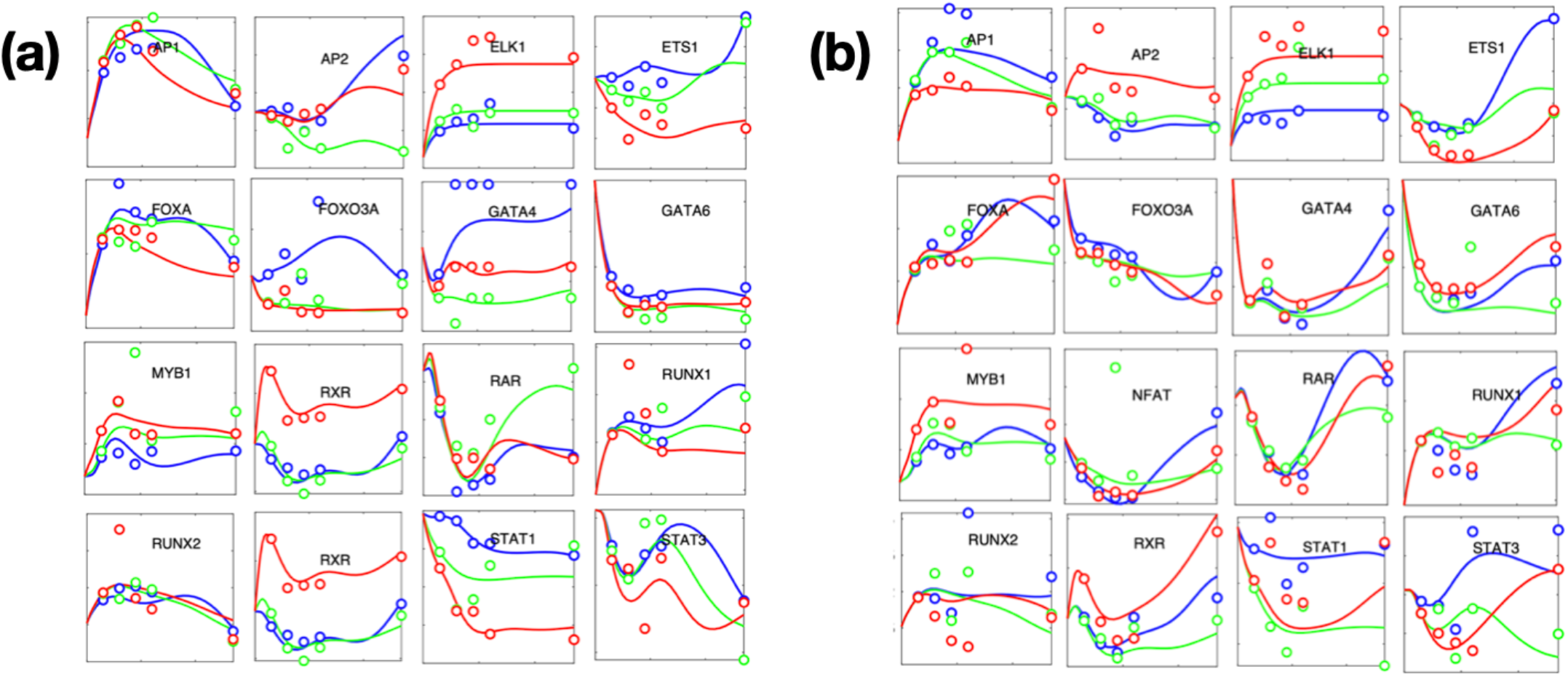
Global optimization for gene regulatory network integration. The subnetworks are integrated by a combination of stochastic global optimization and gradient-based programming (GCMMA) while estimating kinetic parameters from 6 different cues with the gene-to-gene interacting connectivity fixed (Figure 3c). However, the cue-to-gene connectivities are different for each of 8 replicates, and their “overall” connectivities are determined by their statistical significance. (a) Reconstructed gene profiles with 3 RGD peptide levels (low: blue, middle: green, high: red) and a medium level of hydrogel stiffness (b) Reconstructed gene profiles with 3 PEG levels for different hydrogel stiffnesses (soft: blue, medium: green, hard: red) and RGD level in the middle. For (a) and (b), the solid curves are from reconstructed kinetic models, and five circles are from TRACER data (3, 6, 9, 12, 27 hours).

The directly associated genes from RGD and hydrogel stiffness cues are determined statistically from 8 replicates of the hybrid global optimization approach. We assume that statistically significant direct associations exist as activation or inhibition constitutively, i.e. commonly exist for all 6 different external conditions. The *p*-values are calculated from statistical distribution of cue-to-gene association occurrence events through the following steps:

1. For each of 8 replicates, the association from each cue (RGD or hydrogel stiffness) to genes is identified if the input strengths 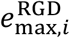 and 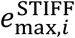 in Equation (2) are ranked within 5 for both activating and inhibiting roles.
2. Regarding cue-to-gene interactions, the occurrences of activation and inhibition of each cue to a specific gene are counted, from which a probability density distribution of occurrences of interaction is obtained. There is a mapping *M*(*i, j*) from each cue *i* to its occurrence of interaction with the specific gene *j*.
3. From these probability density distributions of occurrences, we derive cumulative probability density distributions for activation and inhibition interactions, respectively.
4. For each interaction from cue *i* to gene *j* (activation or inhibition), the occurrence of interaction *M*(*i, j*) points to the corresponding cumulative probability density “*p*_cummulation_”.
5. The cue-to-gene interactions are identified as significant for 1-*p*_cummulation_< 0.05 and colored as green lines (Figure 6).
6. The inferred cue-to-gene interactions are validated with *a priori* known gene databases (Table 1) and identified as validated (solid green lines) when there exists a prior report of the specific cue-to-gene interaction.

In the cue-gene interactions, the inferred activations (Figure 6, green arrows) are still “activation” and “up-regulation” whether the cues are strong or weak. Similarly, the inferred inhibitions (Figure 6, green --|) are still “inhibition” and “down-regulation” independent to the strengths of the cues.

**Figure 6.**
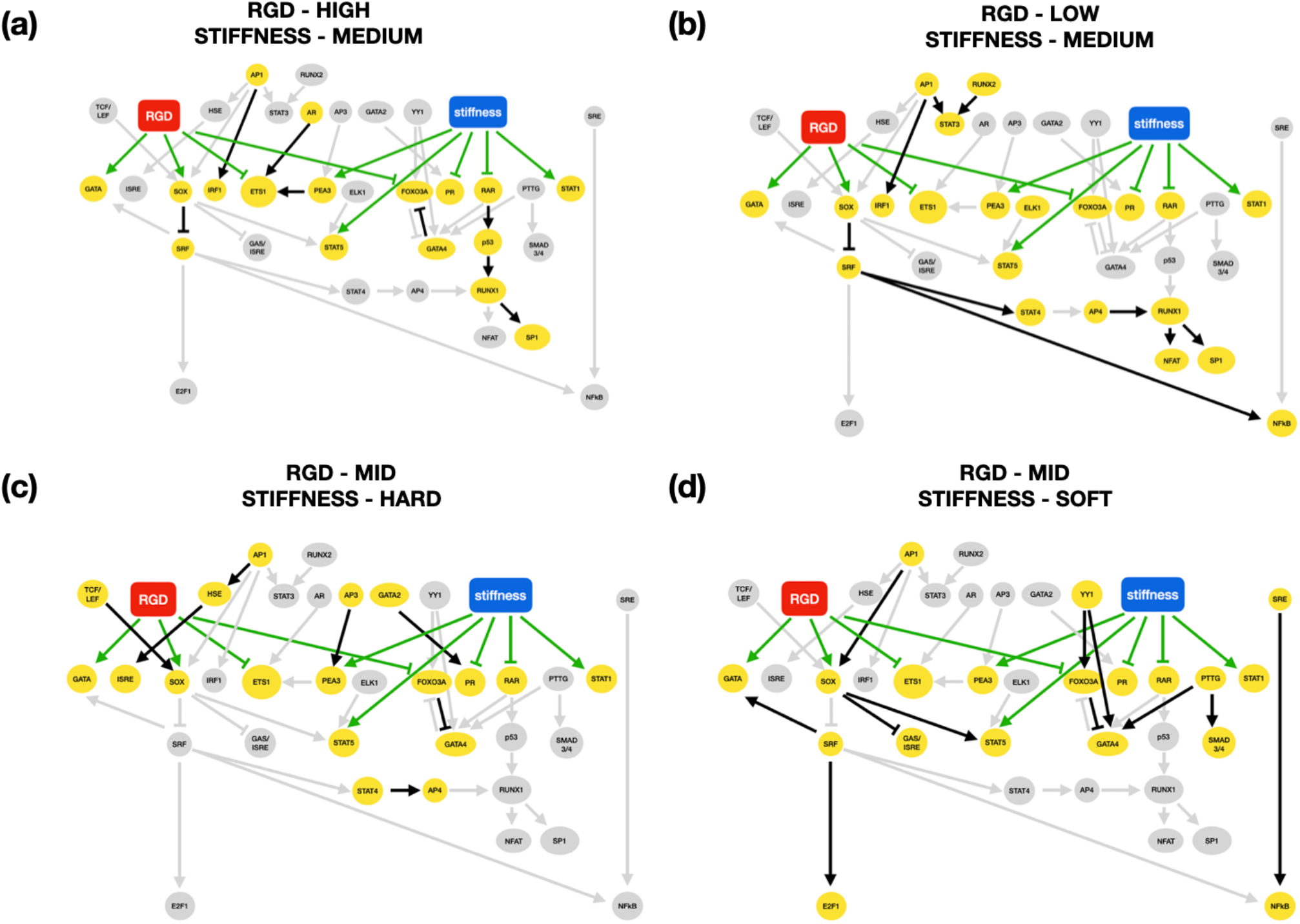
Identified gene network pathways are validated from a priori known gene databases. By overlapping *p*-value based gene-to-gene connectivities commonly significant from two conditions, *differential* networks are constructed. The significant interactions from dual cues to targeted genes are shown by green colored lines (activation → or inhibition --|). They assume to exist constitutively through 6 different conditions. The inferred four networks are validated with *a priori* known gene databases and identified as “*relevant*” (solid) when there exist specific gene-to-gene interactions even though the cell line and conditions may not be the same as those of the TRACER experiments (Table). In the specific conditions, gray colored arrows and genes are latent and only yellow-colored transcriptional factors are effectively involving. (a) Overlapping (RGD-high and stiffness-medium) with (RGD-middle, stiffness-medium). We have inferred that 19 gene-to-gene interactions are relevant among 26 predicted edges. (b) Overlapping (RGD-middle and stiffness-medium) with (RGD-low, stiffness-medium). Among 44 gene-to-gene interactions, 18 are relevant. (c) Overlapping (RGD-middle and stiffness-hard) with (RGD-middle and stiffness-medium). Among 33 gene-to-gene interactions, 18 are relevant. (d) Overlapping (RGD-mid and stiffness-medium) with (RGD-middle and stiffness-soft). Among 56 gene-to-gene interactions, 19 are relevant.

**Table 1:**
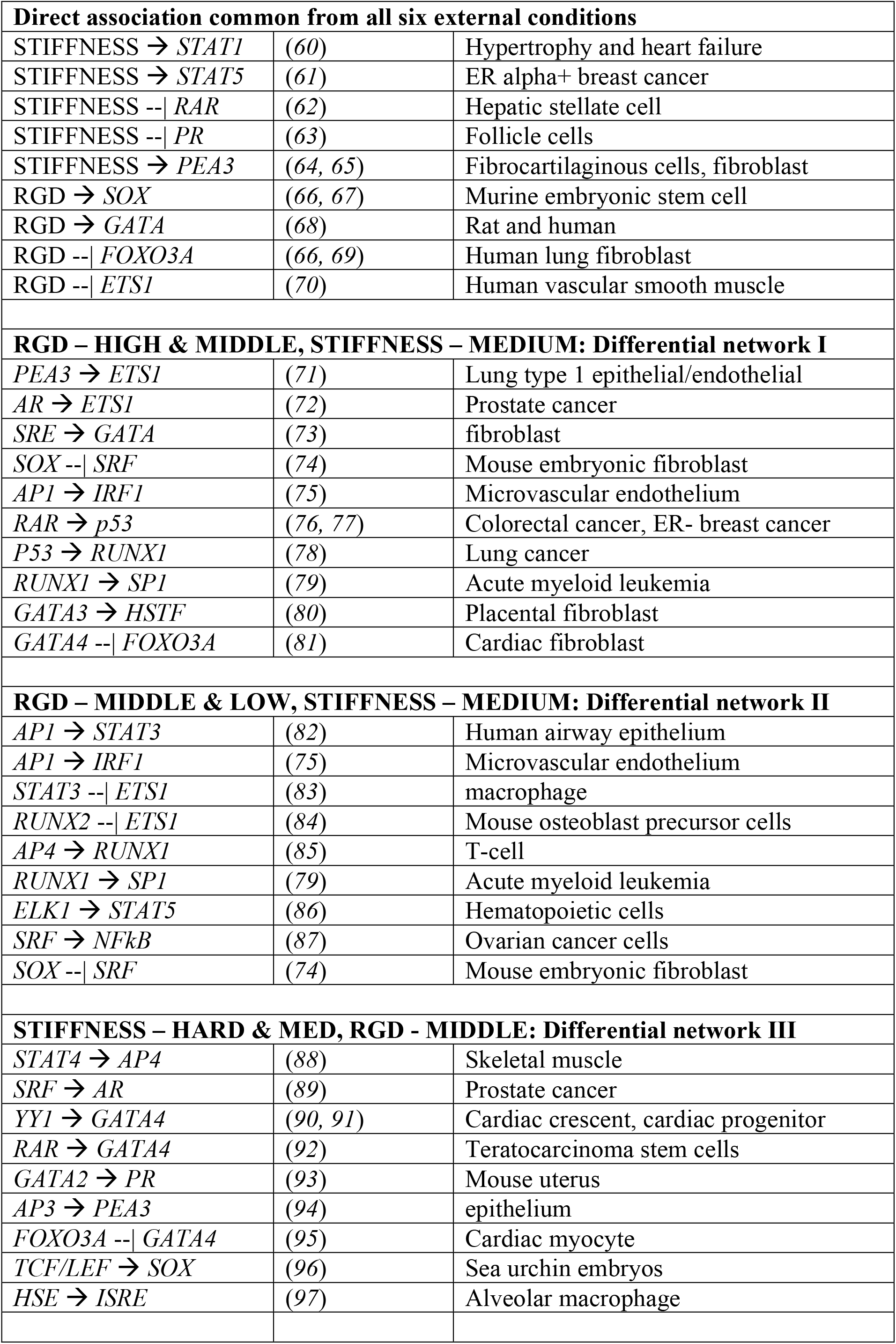
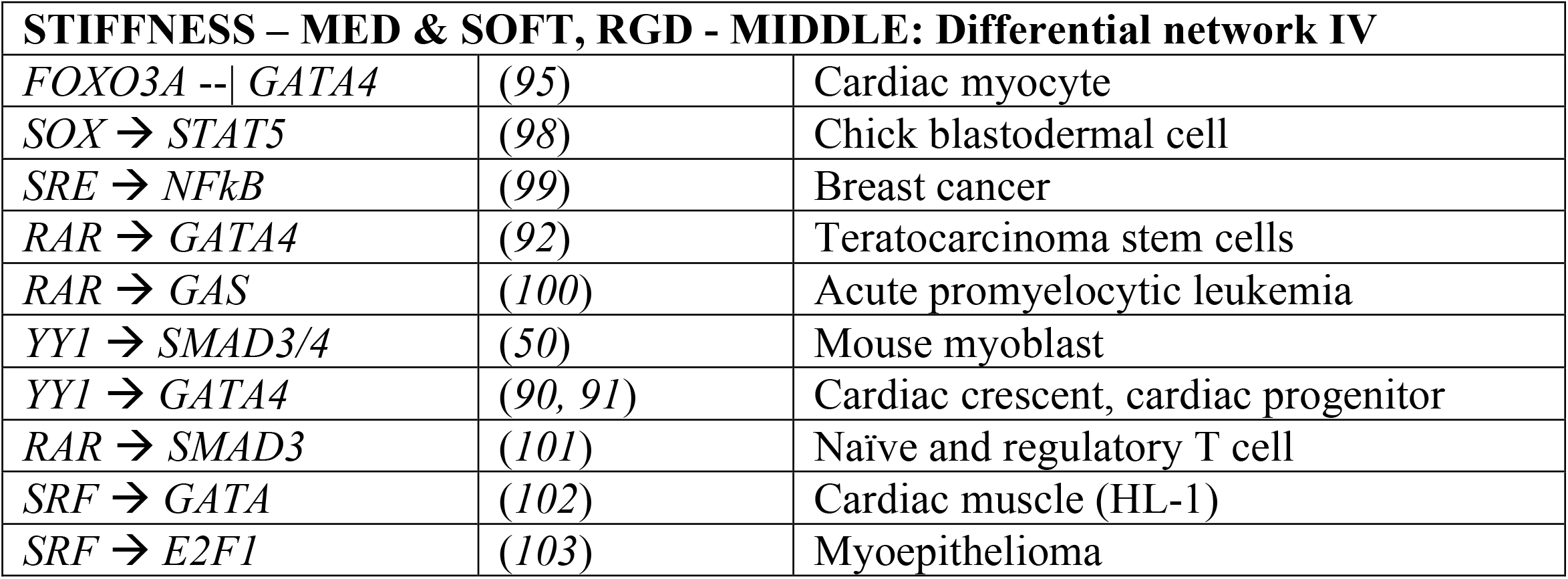
Gene-to-gene interaction validation.

Similarly, statistically significant gene-to-gene interactions are identified for each condition (Figure 4). By overlapping significant (*p*<0.05) gene-to-gene connectivities from two conditions (e.g., RGD-high and RGD-middle, RGD-middle and RGD-low), differential networks are constructed (Figure 6, black colored). The inferred four networks are validated with *a priori* known gene databases (Table 1) and identified as validated (solid lines) when there exists a prior report of the specific gene-to-gene interaction.

## Results and Discussion

### Identified gene network pathways are validated from a priori known gene databases

As detailed in Table 1, of 159 total interactions identified in our global analysis, 74 are validated based on prior reports in the literature.

Hard hydrogel stiffness is predicted to activate *STAT1, STAT5*, and *PEA3*, but inhibit RAR and PR. High level of RGD peptide is predicted to activate stemness-associated *SOX* and *GATA*. However, anti-hypertrophic transcriptional factor *FOXO3A* and ECM remodeling associated *ETS-1* are predicted to be inhibited by high level of RGD peptide. We have identified these direct associations of mechano-transduction as common through in all six experimental conditions. In other words, these upstream interactions from dual cues (cell-ECM adhesion and ECM stiffness) to directly associated genes are prescribed are predicted to be active for all differential regulatory networks (Figure 6, green lines). The identified four differential networks have the following features:

#### (Differential network-I) Overlapping significant gene-to-gene interactions from (RGD-high and hydrogel stiffness-medium) with those from (RGD-middle and hydrogel stiffness-medium)

Among 26 gene-to-gene interactions, 19 are predicted to be active under these conditions (Figure 6a). Several of these inferred pathways may be validated based on *a priori* known databases. When RGD is high, *ETS-1* is inhibited by RGD, but co-activated by *AR* and hydrogel stiffness mediated by *PEA3. RAR* inhibited by ECM stiffness is transmitted through *RAR* → *p53* → *RUNX1* → *SP1. GATA* is coactivated by RGD and *SRE*. At the same time, *FOXO3A* is co-inhibited by *GATA4* and RGD.

#### (Differential network-II) Overlapping significant gene-to-gene interactions from (RGD-middle and hydrogel stiffness-medium) with those from (RGD-low and hydrogel stiffness-medium)

Among 44 gene-to-gene interactions, 18 are predicted to be active under these conditions (Figure 6b). When RGD is low, RGD is inferred to activate *NFkB* through *SRF* (RGD → *SOX* --| *SRF* → *NFkB*). Also, *ETS-1* is co-inhibited by RGD, *RUNX2*, as well as *AP1* mediated by *STAT3*.

#### (Differential network-III) Overlapping significant gene-to-gene interactions from (RGD-middle and hydrogel stiffness-hard) with those from (RGD-middle and hydrogel stiffness-medium)

Among 33 gene-to-gene interactions, 18 are predicted to be active under these conditions (Figure 6c). When hydrogel is stiffness is high, PR is inhibited by stiffness but activated by *GATA2. SOX* is co-activated by *TCF/LEF* and RGD. *GATA4* is activated by *YY1* and RGD mediated by *FOXO3A* but inhibited by hydrogel stiffness mediated by *RAR*.

#### (Differential network-IV) Overlapping significant gene-to-gene interactions from (RGD-mid and hydrogel stiffness-medium) with those from (RGD-mid and hydrogel stiffness-soft)

Among 56 gene-to-gene interactions, 19 are predicted to be active under these conditions (Figure 6d). When hydrogel stiffness is low, *GATA* is co-activated by RGD and *SRF*. Also, *RAR* inhibition is attenuated by soft hydrogel and transmitted to activate *GAS, GATA4*, and *SMAD3/4*.

### Master regulatory transcriptional factors GATA4, SMAD3/4, ETS-1, and STAT5 and associated pathways respond adaptively with differential conditions from mechano-transduction

#### *GATA4* and *SMAD3/4*

As illustrated in Figure 7a, *GATA4* and *SMAD3/4* are differentially regulated based on relative levels of RGD and hydrogel stiffness. The pro-fibrotic factor *SMAD3/4* is inhibited under medium and high hydrogel stiffness conditions, via inhibition of *RAR*. High stiffness is also predicted to inhibit the RAR-mediated activation of *GATA4*. However, independent of the stiffness level, *GATA4* can be upregulated by *RGD* at med and high levels via inhibition of *FOX3A* and activation by *YY1*. However, when RGD is high, *GATA4* does “inhibit” *FOXO3A* together with RGD, i.e., *GATA4* - *FOXO3A* has a mutual feedback structure depending on conditions (Figure 7a).

**Figure 7.**
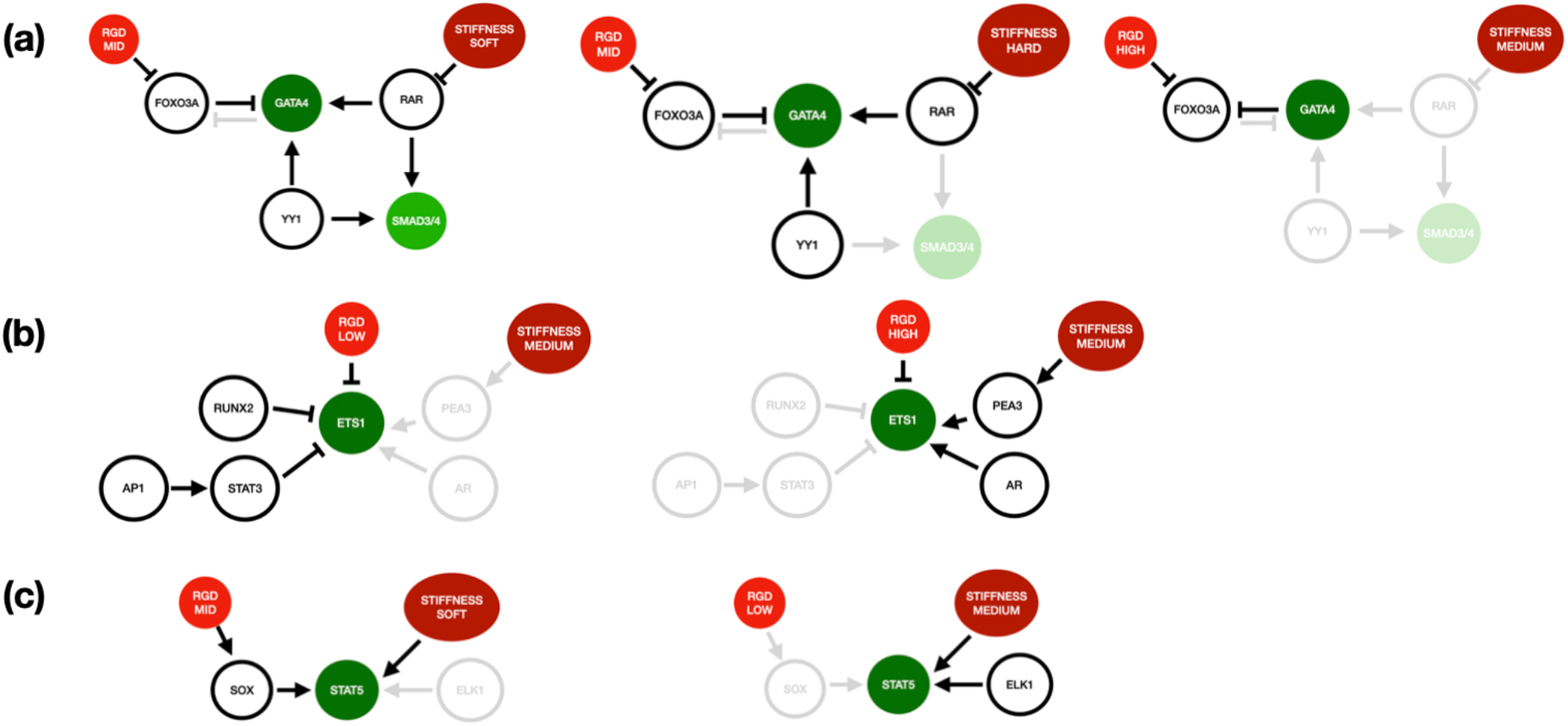
Master regulatory transcriptional factors GATA4, SMAD3/4, ETS-1, and STAT5 and associated pathways respond adaptively with differential conditions from mechano-transduction. Master regulatory genes are identified from validated pathways in Figure 6. The dimmed interactions are not significant in the specific conditions. (a) To indicate the adaptive gene regulations with *GATA4*, the interactions with soft hydrogel stiffness and RGD-middle, those with hard hydrogel stiffness and RGD-middle, those with high RGD and medium hydrogel stiffness are shown in sequence. *GATA4* is counter-stimulated by RGD and hydrogel stiffness, i.e., effectively activated by RGD and inhibited by hydrogel stiffness. Activation of *SMAD3/4* from *YY1* and *RAR* are turned on in soft hydrogel. When RGD is high, *GATA4* inhibits *FOXO3A*. (b) To indicate the adaptive gene regulations with *ETS1*, the interactions with low RGD and medium hydrogel stiffness, and those with high RGD and medium hydrogel stiffness are shown in sequence. *ETS1* is counter-stimulated by RGD and hydrogel stiffness. RGD – low versus high. *ETS1* is inhibited from *STAT3* and *AP1* mediated through *RUNX2* when RGD is low, but activated from medium stiffness mediated by *PEA3* when RGD is high. (c) To indicate the adaptive gene regulations with *STAT5*, the interactions with soft hydrogel stiffness and RGD-middle, and those with low RGD and medium hydrogel stiffness are shown in sequence. *STAT5* is co-stimulated by stiffness and RGD. (RGD-low, stiffness-medium) versus (RGD-mid, stiffness-soft). STAT5 is activated by *ELK1* when RGD is low, but this activator is switched over to *SOX* triggered by the middle level of RGD and soft hydrogel.

#### ETS-1

The target gene *ETS-1* is also predicted to switch expression level depending on the RGD level and hydrogel stiffness, as illustrated in Figure 7b. When the RGD level is low, *AP1* and *RUNX2* co-inhibit *ETS-1* together with RGD. However, when the RGD level is high, *AR* and hydrogel stiffness mediated by *PEA3* “co-activate” *ETS-1*. The stability of *ETS-1* is adaptively regulated by the RGD level and hydrogel stiffness (Figure 7b).

#### STAT5

*STAT5* is a potential inflammatory signal. As illustrated in Figure 7c, STAT5 may be directly activated by either RGD for hydrogel stiffness. When RGD is low, medium or high hydrogel stiffness is required to activate *STAT5* expression (which is also stimulated by *ELK1*). For medium and high levels of *RGD, STAT5* activation by SOX is able to overcome low hydrogel stiffness.

### Prediction of specific phenotypes

#### (Phenotype I) Fibrotic transdifferentiation

TGF-ß and *SMAD3/4* (*49*) are primary players in the fibrotic transdifferentiation from fibroblast to myofibroblast (*50*). Our analysis predicts that mechano-transduction, i.e., when the hydrogel is soft, and *RAR* does mediate, shows a fibrotic potential with targeting *SMAD3/4*.

#### (Phenotype II) Regulation of inflammation and hypertrophy

Fibroblast deeply involves in hypertrophic and inflammatory phenotypes. Our analysis predicts those potentials by *GATA4* (*51-53*) and *STAT5* (*54*), respectively. Especially, hard hydrogel and low RGD drive anti-hypertrophic phenotype targeting to *GATA4*.

#### (Phenotype III) Stem cell lineage determination

Fibroblast is a significant component in the extracellular matrix and microenvironment determining differentiation fates in stem cell lineages. Among others, Stiffness --| *RAR* → *p53* → *RUNX1* axis and RGD → *SOX* → *STAT5* might be new pathways to be investigated further. *GATA* also involves in stemness. When RGD is high, *GATA* is co-activated by RGD and *SRE*. When hydrogel is soft, *GATA* is co-activated by RGD and *SRF*.

#### (Phenotype IV) Activation for invasive and metastatic states

In cancer metastasis, the transition to invasive migration is preconditioned by ECM remodeling and protease by MMP proteins. We have identified *ETS-1* as a potential master regulator through mechano-transduction (*55, 56*). When RGD is low, *ETS-1* is inhibited by proliferation driven by *AP1* and *STAT3* and keeps a stationary stage rather than invasive migration. On the other hand, when RGD is high, *ETS-1* is activated by hydrogel stiffness, highly good condition for a metastatic transition.

### Utilization of a priori known pathways as a constraint in subnetwork reconstruction

The present reconstruction process is unbiased and does not utilize *a priori* known pathways. This kind of approach has pros and cons. Through the solely randomized inferences, we may discover a new pathway and functional architecture. This unbiased approach may predict an actual possibility of pathways for phenotypes of rare diseases. However, this does provide higher false discovery rates (FDR) for highly confirmed gene regulatory subnetwork modules. In a compromise, we plan to incorporate highly probable building blocks of pathways as given without randomized filtration depending on the features of transcriptomics and cell types.

### Oscillation in long-run dynamics and identification of coupled oscillators

In the way of a broader investigation of the inferred gene regulatory networks, we have also obtained a long-run dynamics prescribed with multiple conditions. Interestingly several replicates show oscillatory and non-oscillatory responses depending on the conditions, i.e., bifurcating mechano-sensing features. Those oscillatory features imply some sort of coupled oscillators in the deconvoluted networks. Our inferences are based on a period of 27 hours, and this finite data for the reconstruction may provide limited constraints for the real system. For further validating the authenticity for the mechano-transduction induced oscillation, more data in long-run dynamics will clarify this issue. However, some groups have reported the intrinsic subcellular transports between cytoplasm and nucleus in a feedback loop and associated coupled oscillators (*57, 58*).

### Global optimization techniques

We have utilized a hybrid approach combining global stochastic optimization such as simulated annealing and genetic algorithms and local gradient-based nonlinear programming. Some groups have incorporated Particle Swarm Optimization (PSO) (*59*), for which we may test in the future. As nonlinear programming, we have employed a globally convergent method of GCMMA and prescribed a few constraints in fitting the data. Those inequality constraints can be relaxed or further imposed and fine-tuned in the range of the constraints. In the outer loop, we may improve the fitting by a few more iterations of the global-local hybrid optimization.

### Validation of gene network inference by knockdown and knockout

We have proposed the plasticity of fibroblast in mechano-transduction based on the differentially identified pathways depending on the differential conditions. The experimental validation of these predictions is expected to be done by RNAi and CRISPR techniques, and the protein level western blot will confirm the phenotypic functional significance.

## Notes

### Competing Interest Statement

The authors have declared no competing interest.

## References

1. D. E. Discher, P. Janmey, Y. L. Wang, Tissue cells feel and respond to the stiffness of their substrate. Science 310, 1139–1143 (2005).

2. A. J. Engler, S. Sen, H. L. Sweeney, D.E. Discher, Matrix elasticity directs stem cell lineage specification. Cell 126, 677–689 (2006).

3. M. Chiquet, L. Gelman, R. Lutz, S. Maier, From mechanotransduction to extracellular matrix gene expression in fibroblasts. Biochim Biophys Acta 1793, 911–920 (2009).

4. L. K. Sthanam et al., Biophysical regulation of mouse embryonic stem cell fate and genomic integrity by feeder derived matrices. Biomaterials 119, 9–22 (2017).

5. J. T. Connelly et al., Actin and serum response factor transduce physical cues from the microenvironment to regulate epidermal stem cell fate decisions. Nat Cell Biol 12, 711–718 (2010).

6. A. Urciuolo et al., Collagen VI regulates satellite cell self-renewal and muscle regeneration. Nat Commun 4, 1964 (2013).

7. J. W. Shin, D. J. Mooney, Improving Stem Cell Therapeutics with Mechanobiology. Cell Stem Cell 18, 16–19 (2016).

8. P. A. Lalit et al., Lineage Reprogramming of Fibroblasts into Proliferative Induced Cardiac Progenitor Cells by Defined Factors. Cell Stem Cell 18, 354–367 (2016).

9. F. M. Watt, Engineered Microenvironments to Direct Epidermal Stem Cell Behavior at Single-Cell Resolution. Dev Cell 38, 601–609 (2016).

10. M. Notari et al., The local microenvironment limits the regenerative potential of the mouse neonatal heart. Sci Adv 4, eaao5553 (2018).

11. G. R. Cunha, Mesenchymal-epithelial interactions: past, present, and future. Differentiation 76, 578–586 (2008).

12. D. Ribatti, M. Santoiemma, Epithelial-mesenchymal interactions: a fundamental Developmental Biology mechanism. Int J Dev Biol 58, 303–306 (2014).

13. N. O. Enemchukwu et al., Synthetic matrices reveal contributions of ECM biophysical and biochemical properties to epithelial morphogenesis. J Cell Biol 212, 113–124 (2016).

14. J. L. Leight, M. A. Wozniak, S. Chen, M. L. Lynch, C. S. Chen, Matrix rigidity regulates a switch between TGF-β1-induced apoptosis and epithelial-mesenchymal transition. Mol Biol Cell 23, 781–791 (2012).

15. R. M. Carew, B. Wang, P. Kantharidis, The role of EMT in renal fibrosis. Cell Tissue Res 347, 103–116 (2012).

16. R. Du et al., Hypoxia-induced Bmi1 promotes renal tubular epithelial cell-mesenchymal transition and renal fibrosis via PI3K/Akt signal. Mol Biol Cell 25, 2650–2659 (2014).

17. I. Pastushenko et al., Identification of the tumour transition states occurring during EMT. Nature 556, 463–468 (2018).

18. T. Rozario, D. W. DeSimone, The extracellular matrix in development and morphogenesis: a dynamic view. Dev Biol 341, 126–140 (2010).

19. N. A. Bhowmick, E. G. Neilson, H. L. Moses, Stromal fibroblasts in cancer initiation and progression. Nature 432, 332–337 (2004).

20. A. Orimo et al., Stromal fibroblasts present in invasive human breast carcinomas promote tumor growth and angiogenesis through elevated SDF-1/CXCL12 secretion. Cell 121, 335–348 (2005).

21. R. Kalluri, M. Zeisberg, Fibroblasts in cancer. Nat Rev Cancer 6, 392–401 (2006).

22. K. Pietras, A. Ostman, Hallmarks of cancer: interactions with the tumor stroma. Exp Cell Res 316, 1324–1331 (2010).

23. P. Lu, V. M. Weaver, Z. Werb, The extracellular matrix: a dynamic niche in cancer progression. J Cell Biol 196, 395–406 (2012).

24. V. M. Weaver et al., Reversion of the malignant phenotype of human breast cells in three-dimensional culture and in vivo by integrin blocking antibodies. J Cell Biol 137, 231–245 (1997).

25. A. Siletz, E. Kniazeva, J. S. Jeruss, L. D. Shea, Transcription factor networks in invasion-promoting breast carcinoma-associated fibroblasts. Cancer Microenviron 6, 91–107 (2013).

26. A. De Boeck et al., Differential secretome analysis of cancer-associated fibroblasts and bone marrow-derived precursors to identify microenvironmental regulators of colon cancer progression. Proteomics 13, 379–388 (2013).

27. A. Naba et al., Extracellular matrix signatures of human primary metastatic colon cancers and their metastases to liver. BMC Cancer 14, 518 (2014).

28. L. E. Barney et al., A cell-ECM screening method to predict breast cancer metastasis. Integr Biol (Camb) 7, 198–212 (2015).

29. L. Alba-Castellón et al., Snail1-Dependent Activation of Cancer-Associated Fibroblast Controls Epithelial Tumor Cell Invasion and Metastasis. Cancer Res 76, 6205–6217 (2016).

30. B. Y. Nabet et al., Exosome RNA Unshielding Couples Stromal Activation to Pattern Recognition Receptor Signaling in Cancer. Cell 170, 352-366.e313 (2017).

31. A. Costa et al., Fibroblast Heterogeneity and Immunosuppressive Environment in Human Breast Cancer. Cancer Cell 33, 463-479.e410 (2018).

32. A. J. Booth et al., Acellular normal and fibrotic human lung matrices as a culture system for in vitro investigation. Am J Respir Crit Care Med 186, 866–876 (2012).

33. C. E. Barkauskas, P. W. Noble, Cellular mechanisms of tissue fibrosis. 7. New insights into the cellular mechanisms of pulmonary fibrosis. Am J Physiol Cell Physiol 306, C987–996 (2014).

34. A. Gabrielli, E. V. Avvedimento, T. Krieg, Scleroderma. N Engl J Med 360, 1989–2003 (2009).

35. T. R. Katsumoto, M. L. Whitfield, M. K. Connolly, The pathogenesis of systemic sclerosis. Annu Rev Pathol 6, 509–537 (2011).

36. B. Penalver Bernabe et al., Dynamic transcription factor activity networks in response to independently altered mechanical and adhesive microenvironmental cues. Integrative biology:quantitative biosciences from nano to macro 8, 844–860 (2016).

37. N. Friedman, Inferring cellular networks using probabilistic graphical models. Science 303, 799–805 (2004).

38. A. A. Margolin et al., ARACNE: an algorithm for the reconstruction of gene regulatory networks in a mammalian cellular context. BMC Bioinformatics 7 Suppl 1, S7 (2006).

39. V. A. Huynh-Thu, A. Irrthum, L. Wehenkel, P. Geurts, Inferring regulatory networks from expression data using tree-based methods. PLoS One 5, (2010).

40. N. V. Xuan, M. Chetty, R. Coppel, P. P. Wangikar, Gene regulatory network modeling via global optimization of high-order dynamic Bayesian network. BMC Bioinformatics 13, 131 (2012).

41. A. C. Haury, F. Mordelet, P. Vera-Licona, J. P. Vert, TIGRESS: Trustful Inference of Gene REgulation using Stability Selection. BMC Syst Biol 6, 145 (2012).

42. C. Trapnell et al., The dynamics and regulators of cell fate decisions are revealed by pseudotemporal ordering of single cells. Nat Biotechnol 32, 381–386 (2014).

43. M. Setty et al., Wishbone identifies bifurcating developmental trajectories from single-cell data. Nat Biotechnol 34, 637–645 (2016).

44. E. Marco et al., Bifurcation analysis of single-cell gene expression data reveals epigenetic landscape. Proc Natl Acad Sci U S A 111, E5643–5650 (2014).

45. L. Haghverdi, M. Büttner, F. A. Wolf, F. Buettner, F.J. Theis, Diffusion pseudotime robustly reconstructs lineage branching. Nat Methods 13, 845–848 (2016).

46. J. N. Bazil, F. Qi, D. A. Beard, A parallel algorithm for reverse engineering of biological networks. Integrative biology:quantitative biosciences from nano to macro 3, 1215–1223 (2011).

47. R. Thiagarajan, A. Alavi, J. T. Podichetty, J. N. Bazil, D. A. Beard, The feasibility of genome-scale biological network inference using Graphics Processing Units. Algorithms Mol Biol 12, 8 (2017).

48. K. Svanberg, A Class of globally convergent optimization methods based on conservative convex separable approximations. SIAM Journal of Optimization 12, 555–573 (2002).

49. J. Massagué, Seoane, J., and Wotton, D., Smad transcription factors. Genes Dev. 19, 2783–2810 (2005).

50. L. Zhou et al., Inhibition of miR-29 by TGF-beta-Smad3 signaling through dual mechanisms promotes transdifferentiation of mouse myoblasts into myofibroblasts. PLoS One 7, e33766 (2012).

51. C. Kang et al., The DNA damage response induces inflammation and senescence by inhibiting autophagy of GATA4. Science 349, aaa5612 (2015).

52. N. Pilon, D. Raiwet, R. S. Viger, D. W. Silversides, Novel pre- and post-gastrulation expression of Gata4 within cells of the inner cell mass and migratory neural crest cells. Dev Dyn 237, 1133–1143 (2008).

53. R. Zheng, G. A. Blobel, GATA Transcription Factors and Cancer. Genes Cancer 1, 1178–1188 (2010).

54. G. Ferbeyre, R. Moriggl, The role of Stat5 transcription factors as tumor suppressors or oncogenes. Biochim Biophys Acta 1815, 104–114 (2011).

55. J. Dittmer, The biology of the Ets1 proto-oncogene. Mol Cancer 2, 29 (2003).

56. P. Haines, G. H. Samuel, H. Cohen, M. Trojanowska, A. M. Bujor, Caveolin-1 is a negative regulator of MMP-1 gene expression in human dermal fibroblasts via inhibition of Erk1/2/Ets1 signaling pathway. J Dermatol Sci 64, 210–216 (2011).

57. S. Sampattavanich et al., Encoding Growth Factor Identity in the Temporal Dynamics of FOXO3 under the Combinatorial Control of ERK and AKT Kinases. Cell Syst 6, 664-678.e669 (2018).

58. A. Hubaud, I. Regev, L. Mahadevan, O. Pourquié, Excitable Dynamics and Yap-Dependent Mechanical Cues Drive the Segmentation Clock. Cell 171, 668-682.e611 (2017).

59. S. Iadevaia, Y. Lu, F. C. Morales, G. B. Mills, P. T. Ram, Identification of optimal drug combinations targeting cellular networks: integrating phospho-proteomics and computational network analysis. Cancer Res 70, 6704–6714 (2010).

60. P. Fischer, D. Hilfiker-Kleiner, Survival pathways in hypertrophy and heart failure: the gp130-STAT3 axis. Basic Res Cardiol 102, 279–297 (2007).

61. C. E. Barcus, P. J. Keely, K. W. Eliceiri, L. A. Schuler, Stiff collagen matrices increase tumorigenic prolactin signaling in breast cancer cells. J Biol Chem 288, 12722–12732 (2013).

62. S. H. Yi, Y. Zhang, D. Tang, L. Zhu, Mechanical force and tensile strain activated hepatic stellate cells and inhibited retinol metabolism. Biotechnol Lett 37, 1141–1152 (2015).

63. E. R. West, M. Xu, T. K. Woodruff, L. D. Shea, Physical properties of alginate hydrogels and their effects on in vitro follicle development. Biomaterials 28, 4439–4448 (2007).

64. S. Kapila, Y. Xie, W. Wang, Induction of MMP-1 (collagenase-1) by relaxin in fibrocartilaginous cells requires both the AP-1 and PEA-3 promoter sites. Orthod Craniofac Res 12, 178–186 (2009).

65. A. Petersen, P. Joly, C. Bergmann, G. Korus, G. N. Duda, The impact of substrate stiffness and mechanical loading on fibroblast-induced scaffold remodeling. Tissue Eng Part A 18, 1804–1817 (2012).

66. O. Hazenbiller, N. A. Duncan, R. J. Krawetz, Reduction of pluripotent gene expression in murine embryonic stem cells exposed to mechanical loading or Cyclo RGD peptide. BMC Cell Biol 18, 32 (2017).

67. J. C. Chang, S. H. Hsu, D. C. Chen, The promotion of chondrogenesis in adipose-derived adult stem cells by an RGD-chimeric protein in 3D alginate culture. Biomaterials 30, 6265–6275 (2009).

68. Y. Katoh, M. Katoh, Identification and characterization of rat Wnt6 and Wnt10a genes in silico. Int J Mol Med 15, 527–531 (2005).

69. R. S. Nho, J. Kahm, beta1-Integrin-collagen interaction suppresses FoxO3a by the coordination of Akt and PP2A. J Biol Chem 285, 14195–14209 (2010).

70. J. H. Yang, W. H. Briggs, P. Libby, R. T. Lee, Small mechanical strains selectively suppress matrix metalloproteinase-1 expression by human vascular smooth muscle cells. J Biol Chem 273, 6550–6555 (1998).

71. H. Kathuria, Y. X. Cao, M. I. Ramirez, M. C. Williams, Transcription of the caveolin-1 gene is differentially regulated in lung type I epithelial and endothelial cell lines. A role for ETS proteins in epithelial cell expression. J Biol Chem 279, 30028–30036 (2004).

72. C. E. Massie et al., New androgen receptor genomic targets show an interaction with the ETS1 transcription factor. EMBO Rep 8, 871–878 (2007).

73. N. S. Belaguli et al., Cardiac tissue enriched factors serum response factor and GATA-4 are mutual coregulators. Mol Cell Biol 20, 7550–7558 (2000).

74. S. Huh et al., Suppression of the ERK-SRF axis facilitates somatic cell reprogramming. Exp Mol Med 50, e448 (2018).

75. F. Wu, K. Tyml, J. X. Wilson, iNOS expression requires NADPH oxidase-dependent redox signaling in microvascular endothelial cells. J Cell Physiol 217, 207–214 (2008).

76. C. C. Applegate, M. A. Lane, Role of retinoids in the prevention and treatment of colorectal cancer. World J Gastrointest Oncol 7, 184–203 (2015).

77. A. Shilkaitis, A. Green, K. Christov, Retinoids induce cellular senescence in breast cancer cells by RAR-β dependent and independent pathways: Potential clinical implications (Review). Int J Oncol 47, 35–42 (2015).

78. J. W. Lee, A. van Wijnen, S. C. Bae, RUNX3 and p53: How Two Tumor Suppressors Cooperate Against Oncogenic Ras? Adv Exp Med Biol 962, 321–332 (2017).

79. R. Pippa et al., MYC-dependent recruitment of RUNX1 and GATA2 on the SET oncogene promoter enhances PP2A inactivation in acute myeloid leukemia. Oncotarget 8, 53989–54003 (2017).

80. I. Mineva, M. Stamenova, W. Gartner, L. Wagner, Expression of the small heat shock protein alphaB-crystallin in term human placenta. Am J Reprod Immunol 60, 440–448 (2008).

81. H. Zhou, M. E. Dickson, M. S. Kim, R. Bassel-Duby, E. N. Olson, Akt1/protein kinase B enhances transcriptional reprogramming of fibroblasts to functional cardiomyocytes. Proc Natl Acad Sci U S A 112, 11864–11869 (2015).

82. X. Liu et al., LPS-induced proinflammatory cytokine expression in human airway epithelial cells and macrophages via NF-κB, STAT3 or AP-1 activation. Mol Med Rep 17, 5484–5491 (2018).

83. H. Zhang et al., STAT3 restrains RANK-and TLR4-mediated signalling by suppressing expression of the E2 ubiquitin-conjugating enzyme Ubc13. Nat Commun 5, 5798 (2014).

84. Y. Zhang et al., Co-stimulation of the bone-related Runx2 P1 promoter in mesenchymal cells by SP1 and ETS transcription factors at polymorphic purine-rich DNA sequences (Y-repeats). J Biol Chem 284, 3125–3135 (2009).

85. T. Egawa, D. R. Littman, Transcription factor AP4 modulates reversible and epigenetic silencing of the Cd4 gene. Proc Natl Acad Sci U S A 108, 14873–14878 (2011).

86. H. Gu, J. C. Pratt, S. J. Burakoff, B. G. Neel, Cloning of p97/Gab2, the major SHP2-binding protein in hematopoietic cells, reveals a novel pathway for cytokine-induced gene activation. Mol Cell 2, 729–740 (1998).

87. Y. Deng et al., A Blockade of IGF Signaling Sensitizes Human Ovarian Cancer Cells to the Anthelmintic Niclosamide-Induced Anti-Proliferative and Anticancer Activities. Cell Physiol Biochem 39, 871–888 (2016).

88. C. Dogra, D. S. Srivastava, A. Kumar, Protein-DNA array-based identification of transcription factor activities differentially regulated in skeletal muscle of normal and dystrophin-deficient mdx mice. Mol Cell Biochem 312, 17–24 (2008).

89. M. Prencipe et al., Relationship between serum response factor and androgen receptor in prostate cancer. Prostate 75, 1704–1717 (2015).

90. K. H. Lee, S. Evans, T. Y. Ruan, A. B. Lassar, SMAD-mediated modulation of YY1 activity regulates the BMP response and cardiac-specific expression of a GATA4/5/6-dependent chick Nkx2.5 enhancer. Development 131, 4709–4723 (2004).

91. S. Gregoire et al., Essential and unexpected role of Yin Yang 1 to promote mesodermal cardiac differentiation. Circ Res 112, 900–910 (2013).

92. D. Su, L. J. Gudas, Retinoic acid receptor gamma activates receptor tyrosine kinase Tie1 gene transcription through transcription factor GATA4 in F9 stem cells. Exp Hematol 36, 624–641 (2008).

93. C. A. Rubel, H. L. Franco, J. W. Jeong, J. P. Lydon, F. J. DeMayo, GATA2 is expressed at critical times in the mouse uterus during pregnancy. Gene Expr Patterns 12, 196–203 (2012).

94. A. L. Façanha, M. C. dos Reis, M. Montero-Lomeli, Structural study of the porcine Na+/H+ exchanger NHE1 gene and its 5’-flanking region. Mol Cell Biochem 210, 91–99 (2000).

95. N. R. Sundaresan et al., Sirt3 blocks the cardiac hypertrophic response by augmenting Foxo3a-dependent antioxidant defense mechanisms in mice. J Clin Invest 119, 2758–2771 (2009).

96. J. D. Kormish, D. Sinner, A. M. Zorn, Interactions between SOX factors and Wnt/beta-catenin signaling in development and disease. Dev Dyn 239, 56–68 (2010).

97. M. S. Harkins, P. L. Moseley, G. K. Iwamoto, Regulation of CD23 in the chronic inflammatory response in asthma: a role for interferon-gamma and heat shock protein 70 in the TH2 environment. Ann Allergy Asthma Immunol 91, 567–574 (2003).

98. S. I. Lee et al., The miR-302 cluster transcriptionally regulated by POUV, SOX and STAT5B controls the undifferentiated state through the post-transcriptional repression of PBX3 and E2F7 in early chick development. Mol Reprod Dev 81, 1103–1114 (2014).

99. M. C. Peeters et al., The adhesion G protein-coupled receptor G2 (ADGRG2/GPR64) constitutively activates SRE and NFκB and is involved in cell adhesion and migration. Cell Signal 27, 2579–2588 (2015).

100. M. Stadler et al., Transcriptional induction of the PML growth suppressor gene by interferons is mediated through an ISRE and a GAS element. Oncogene 11, 2565–2573 (1995).

101. J. Nolting et al., Retinoic acid can enhance conversion of naive into regulatory T cells independently of secreted cytokines. J Exp Med 206, 2131–2139 (2009).

102. A. He, S. W. Kong, Q. Ma, W. T. Pu, Co-occupancy by multiple cardiac transcription factors identifies transcriptional enhancers active in heart. Proc Natl Acad Sci U S A 108, 5632–5637 (2011).

103. M. Urbini et al., Identification of SRF-E2F1 fusion transcript in EWSR-negative myoepithelioma of the soft tissue. Oncotarget 8, 60036–60045 (2017).

