## Supplementary Information for "Plasticity of fibroblast transcriptional response to physical and biochemical cues revealed by dynamic network analysis"

**FIGURE S1. Gene profile clustering, RGD-high and Stiffness-hard**

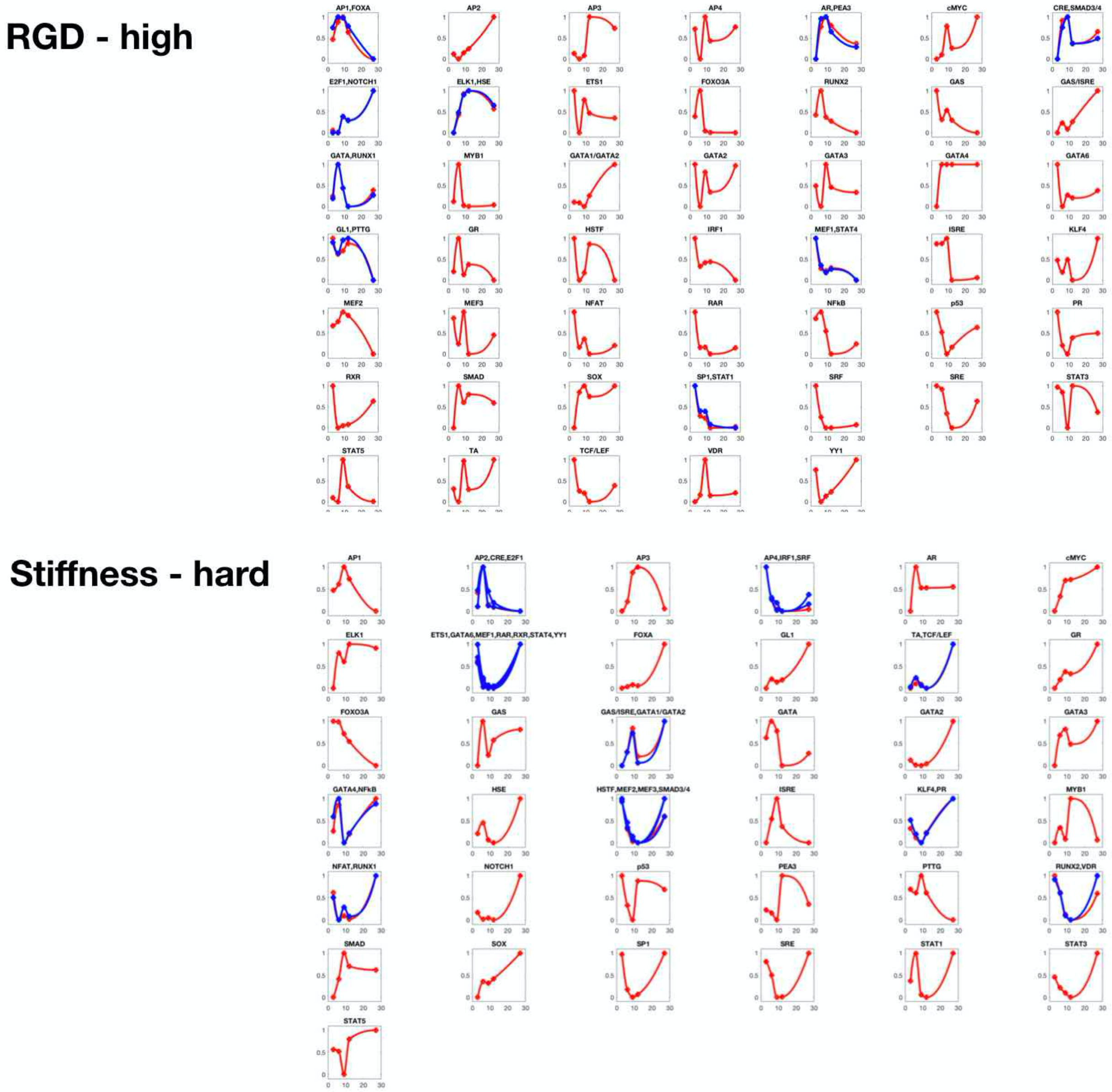

**FIGURE S2. Gene profile clustering, RGD-mid and Stiffness-medium**

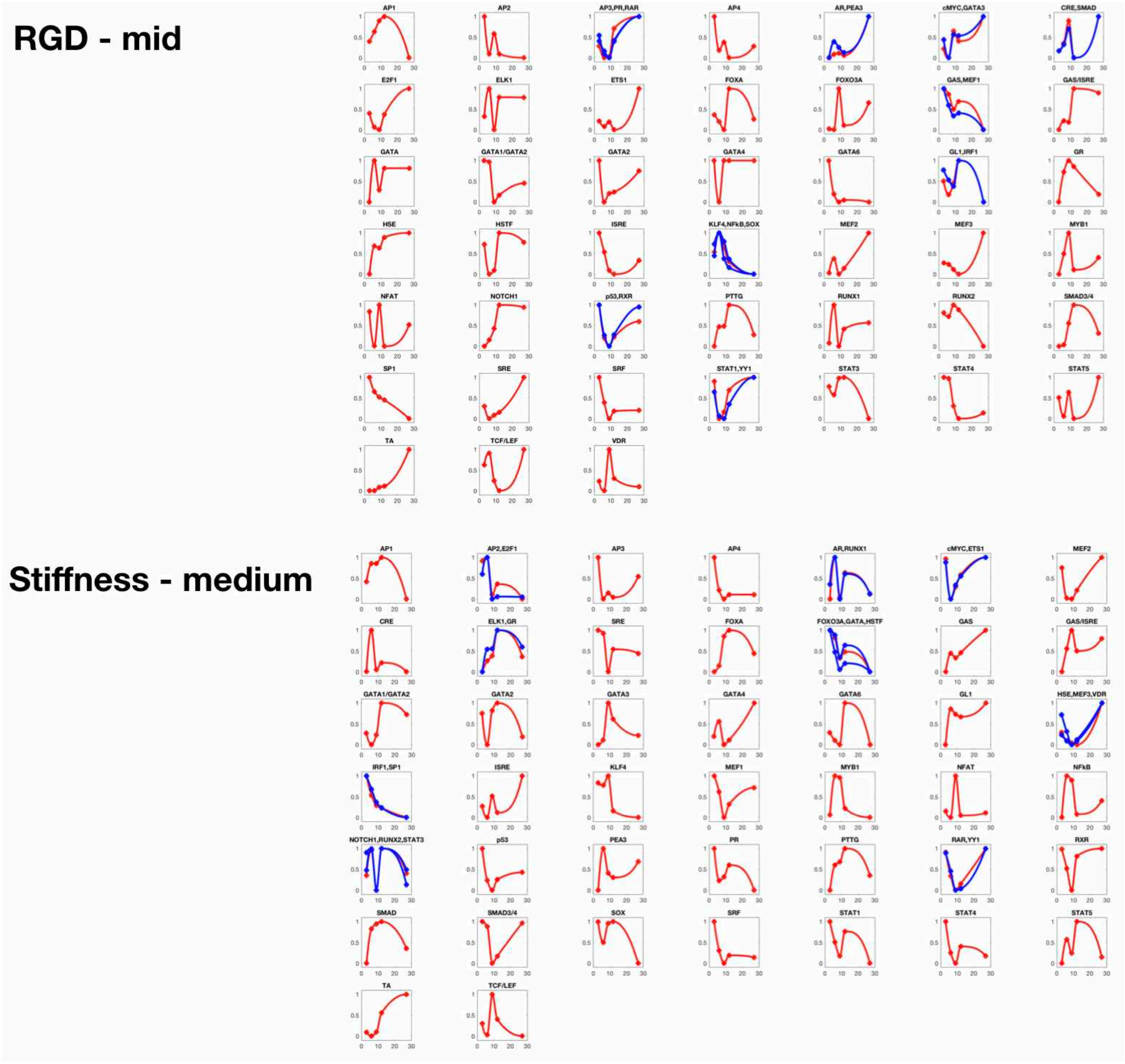

**FIGURE S3. Gene profile clustering, RGD-low and Stiffness-soft**

#### RGD - low

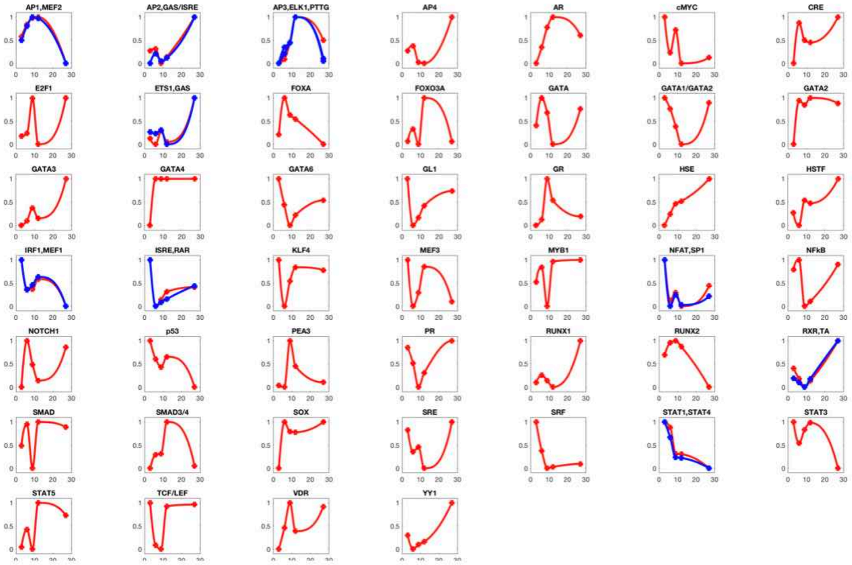

#### Stiffness - soft

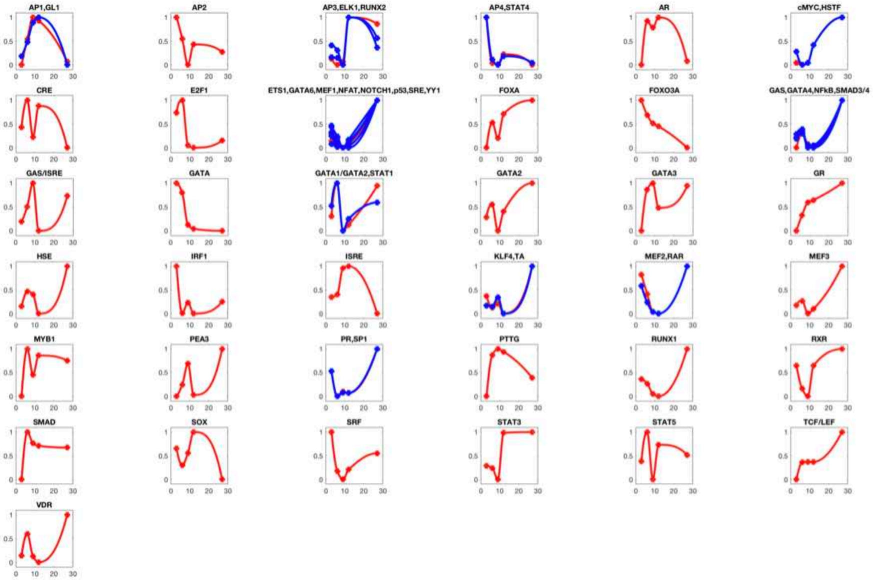

**FIGURE S4. The reconstruction of subnetworks**

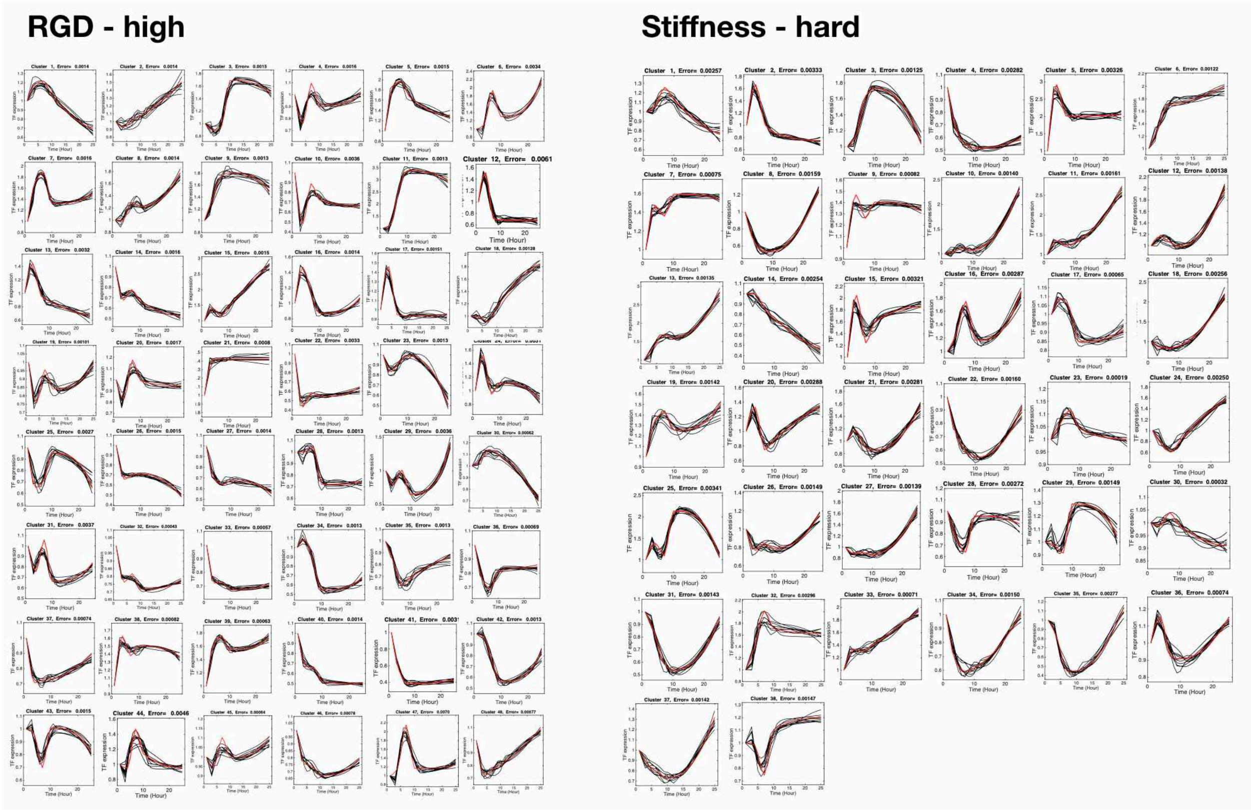


**FIGURE S5. The reconstruction of subnetworks**

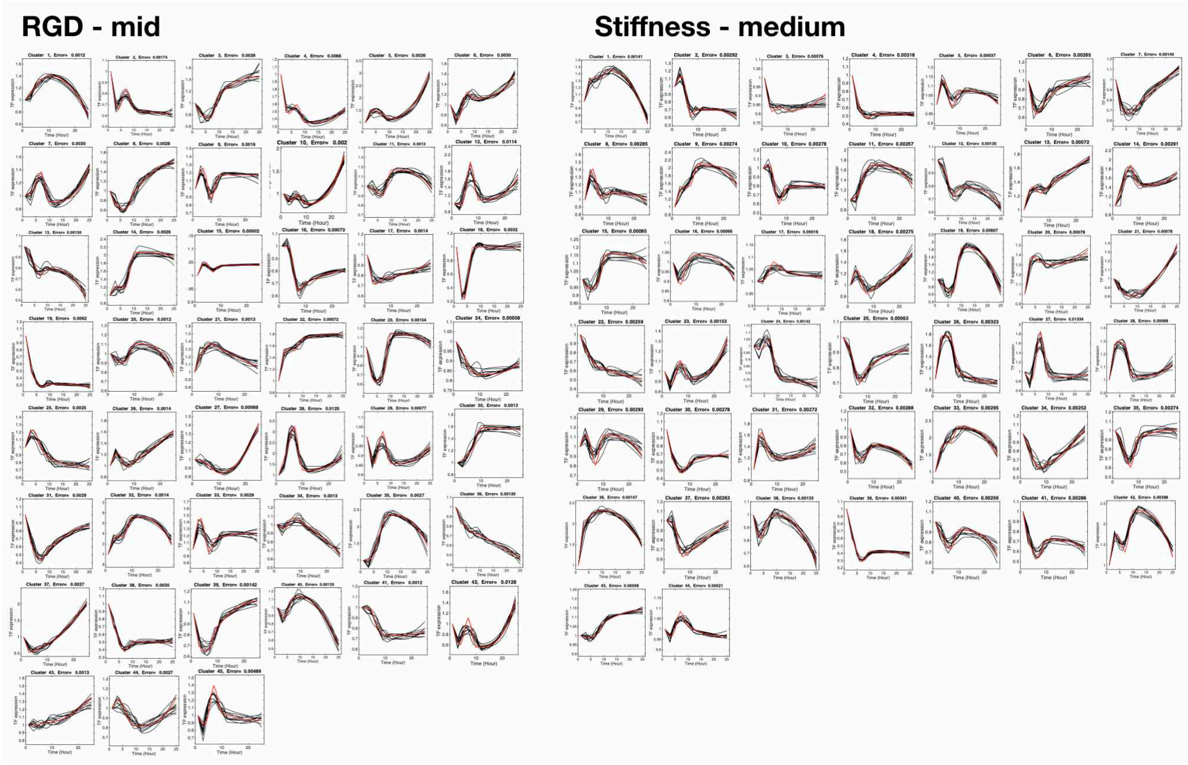

### FIGURE S6. The reconstruction of subnetworks

#### RGD - low

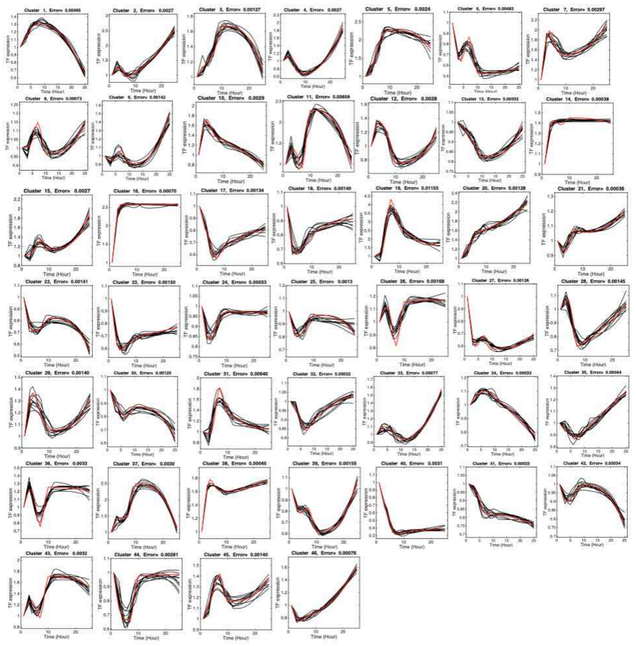

#### Stiffness - soft

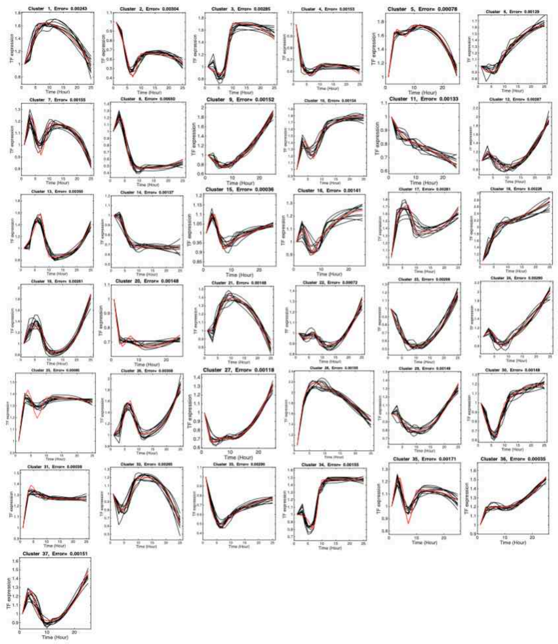

**FIGURE S7. Global gene network optimization**

All integration of subnetworks, i.e. with 6 conditions

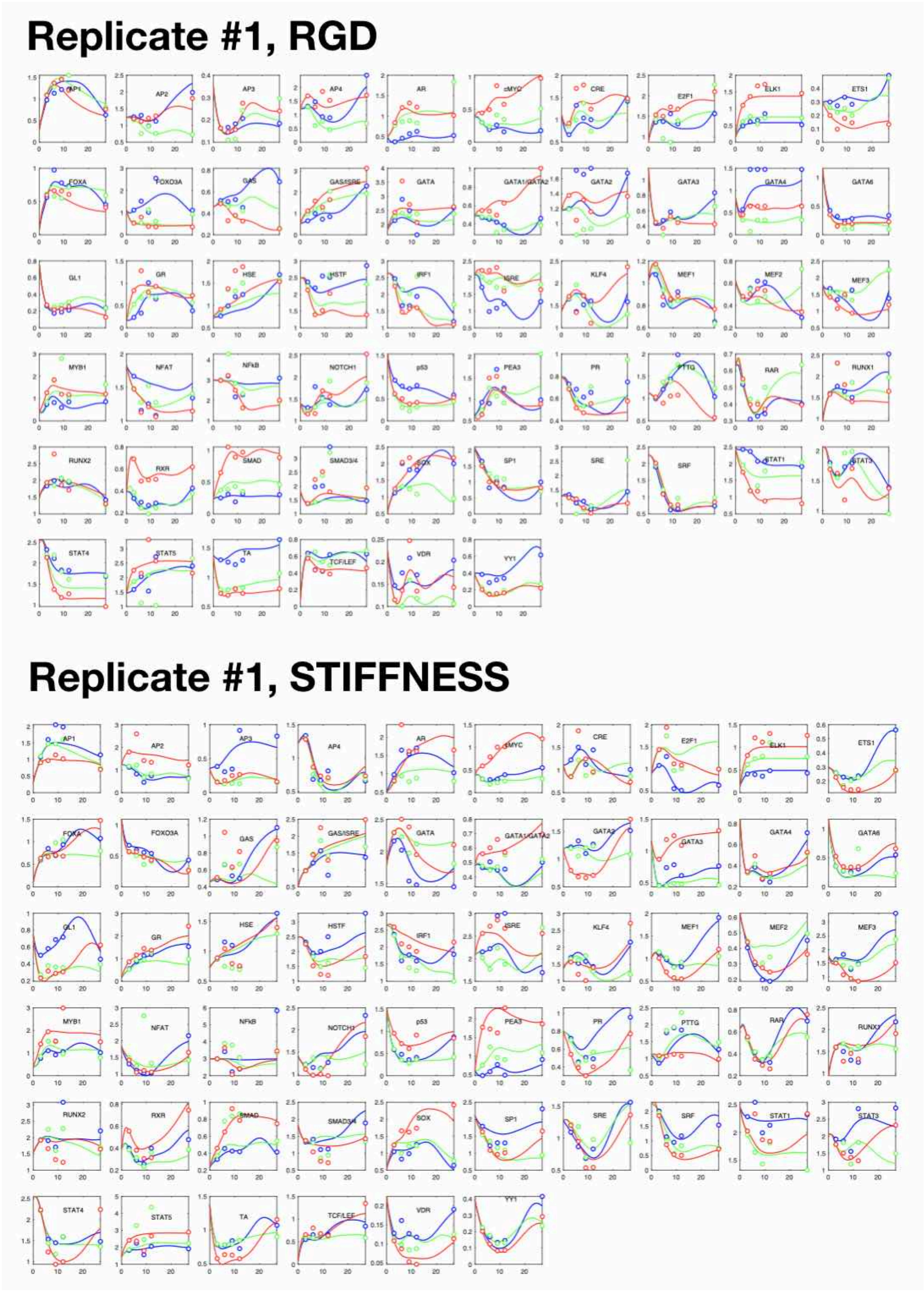

**FIGURE S8. Global gene network optimization**

All integration of subnetworks, i.e. with 6 conditions

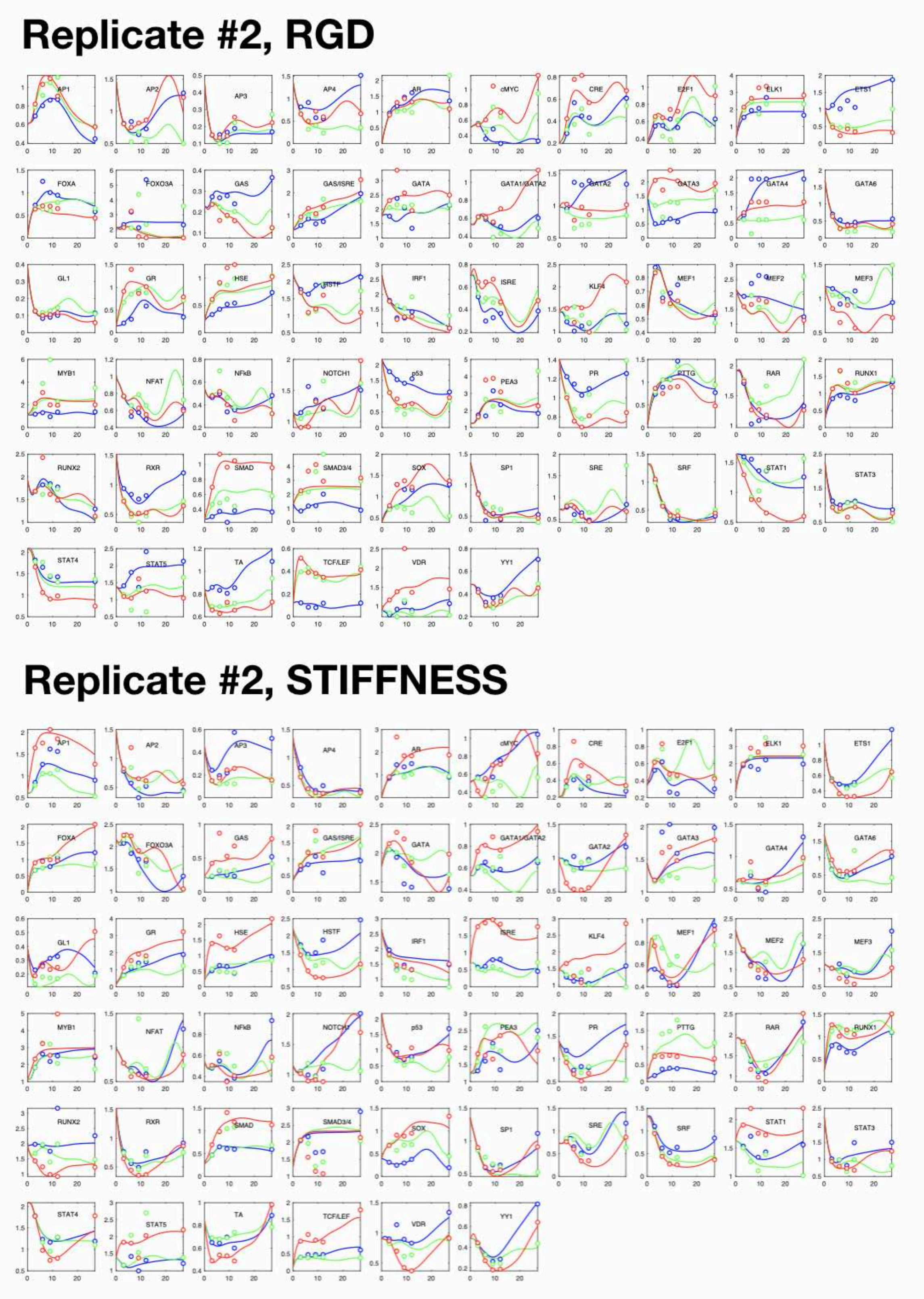

**FIGURE S9. Global gene network optimization**

All integration of subnetworks, i.e. with 6 conditions

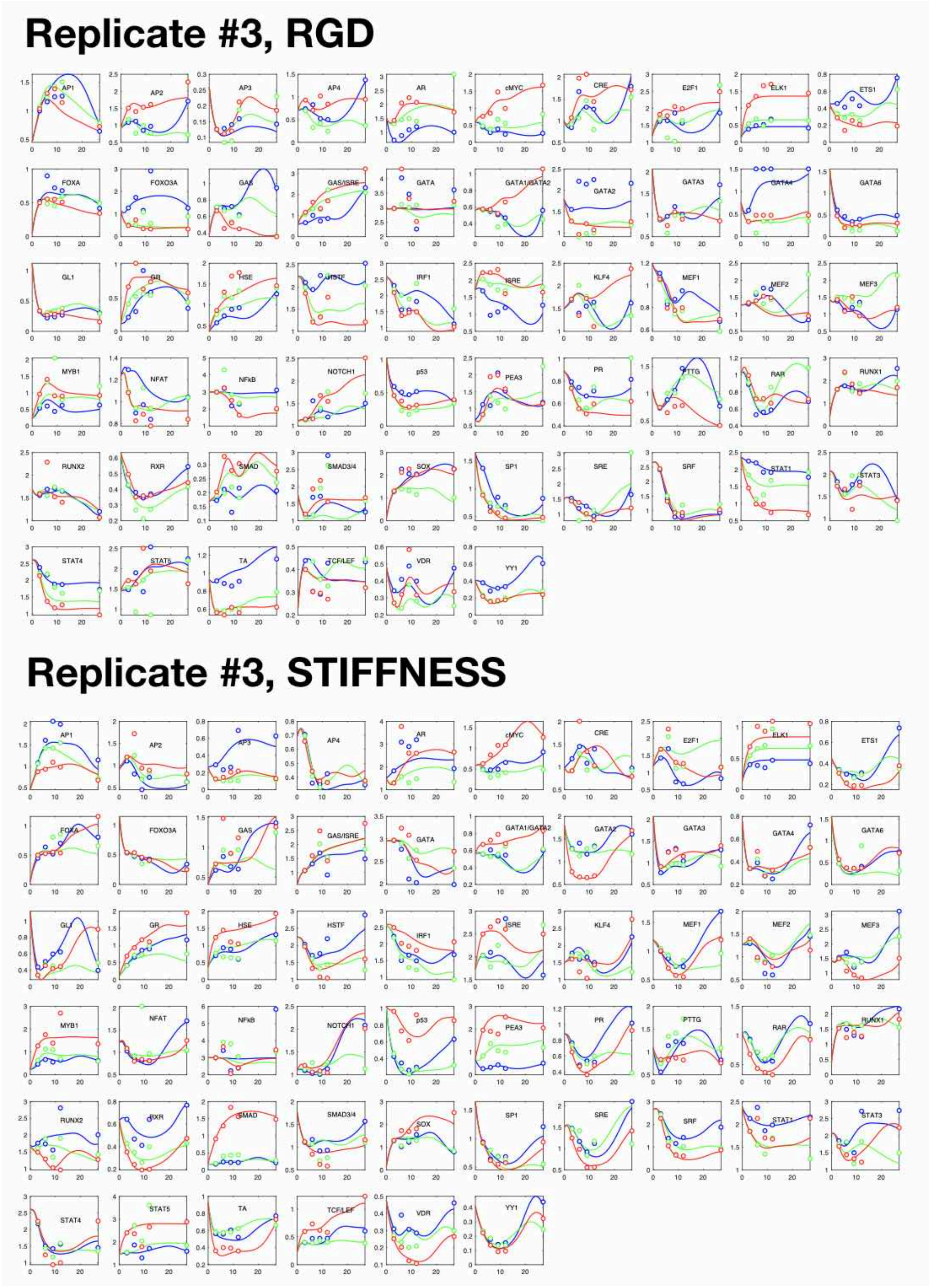

**FIGURE S10. Global gene network optimization**

All integration of subnetworks, i.e. with 6 conditions

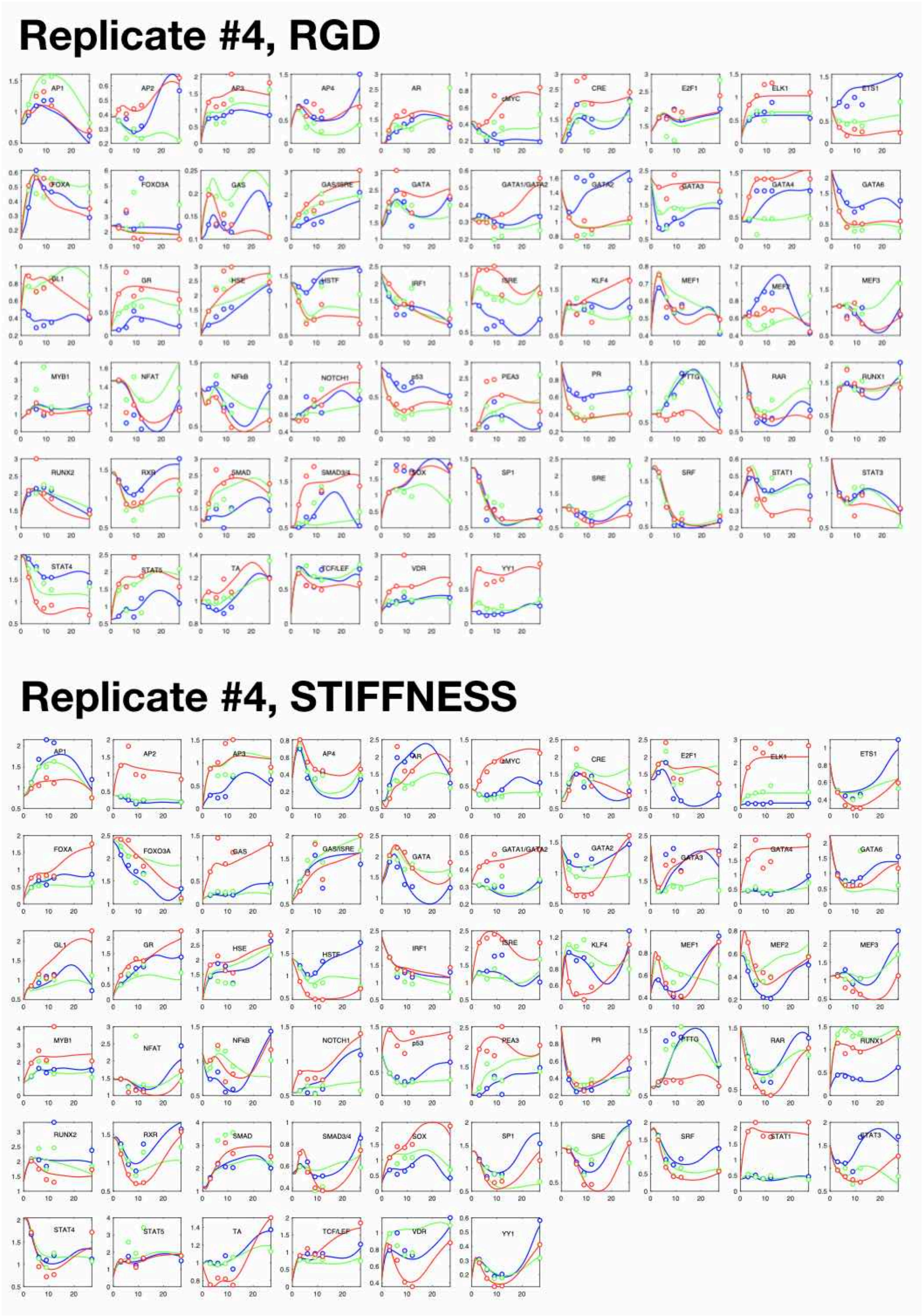

**FIGURE S11. Global gene network optimization**

All integration of subnetworks, i.e. with 6 conditions

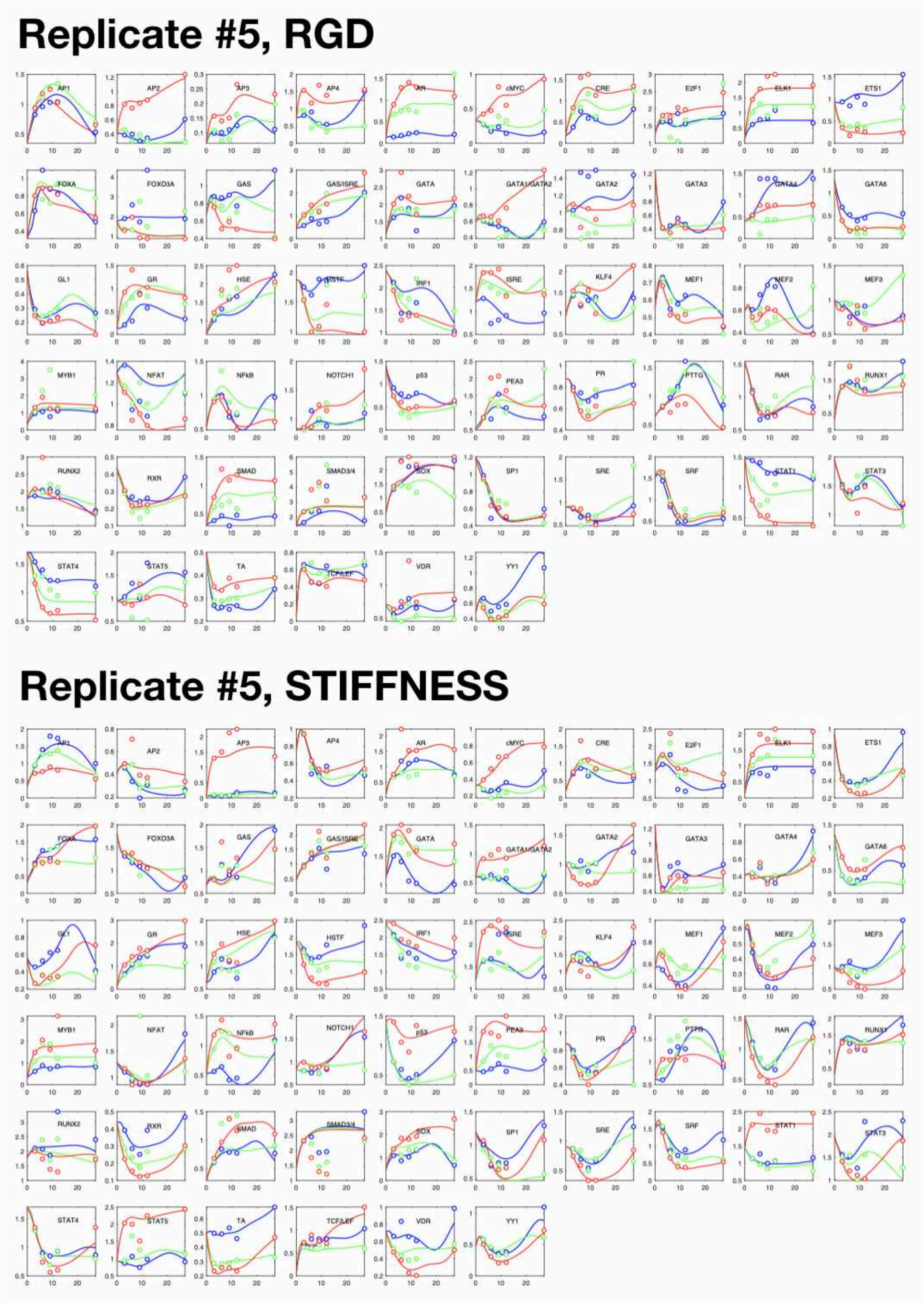

**FIGURE S12. Global gene network optimization**

All integration of subnetworks, i.e. with 6 conditions

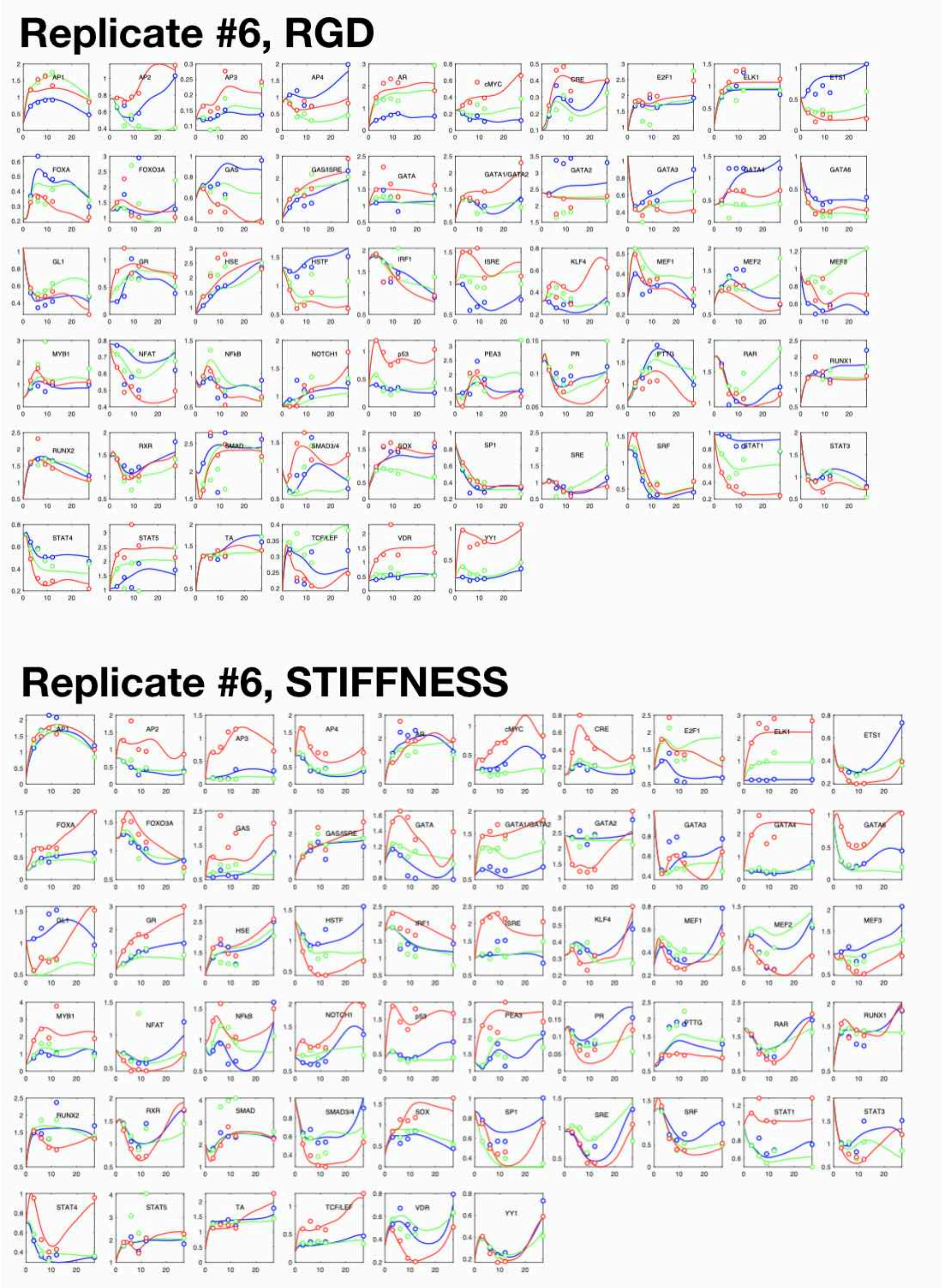

**FIGURE S13. Global gene network optimization**

All integration of subnetworks, i.e. with 6 conditions

#### Replicate #7, RGD

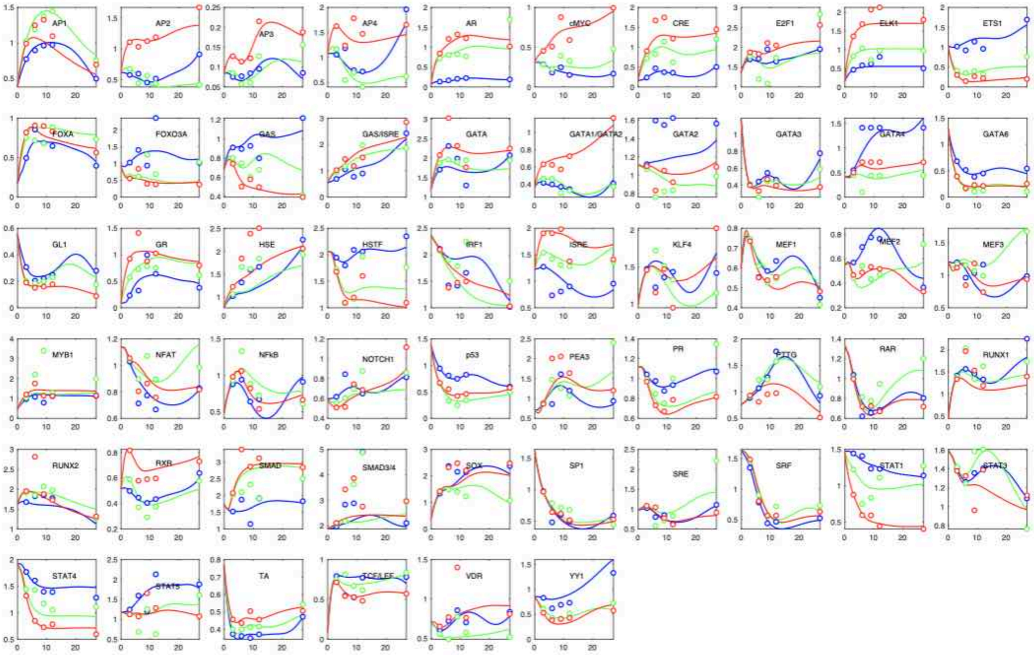


#### Replicate #7, STIFFNESS

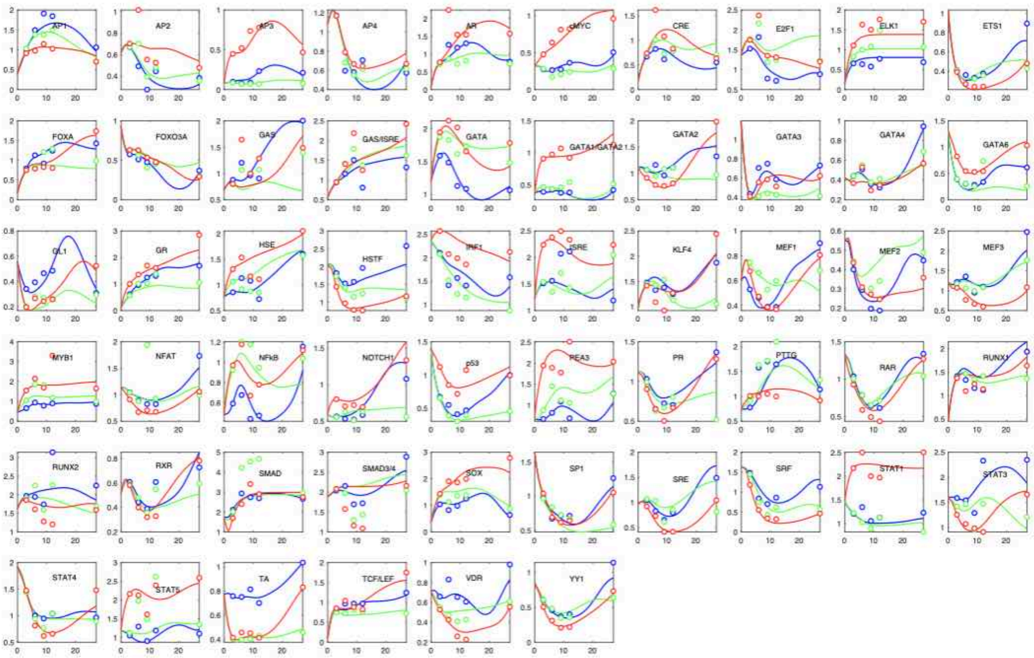

**FIGURE S14. Global gene network optimization**

All integration of subnetworks, i.e. with 6 conditions

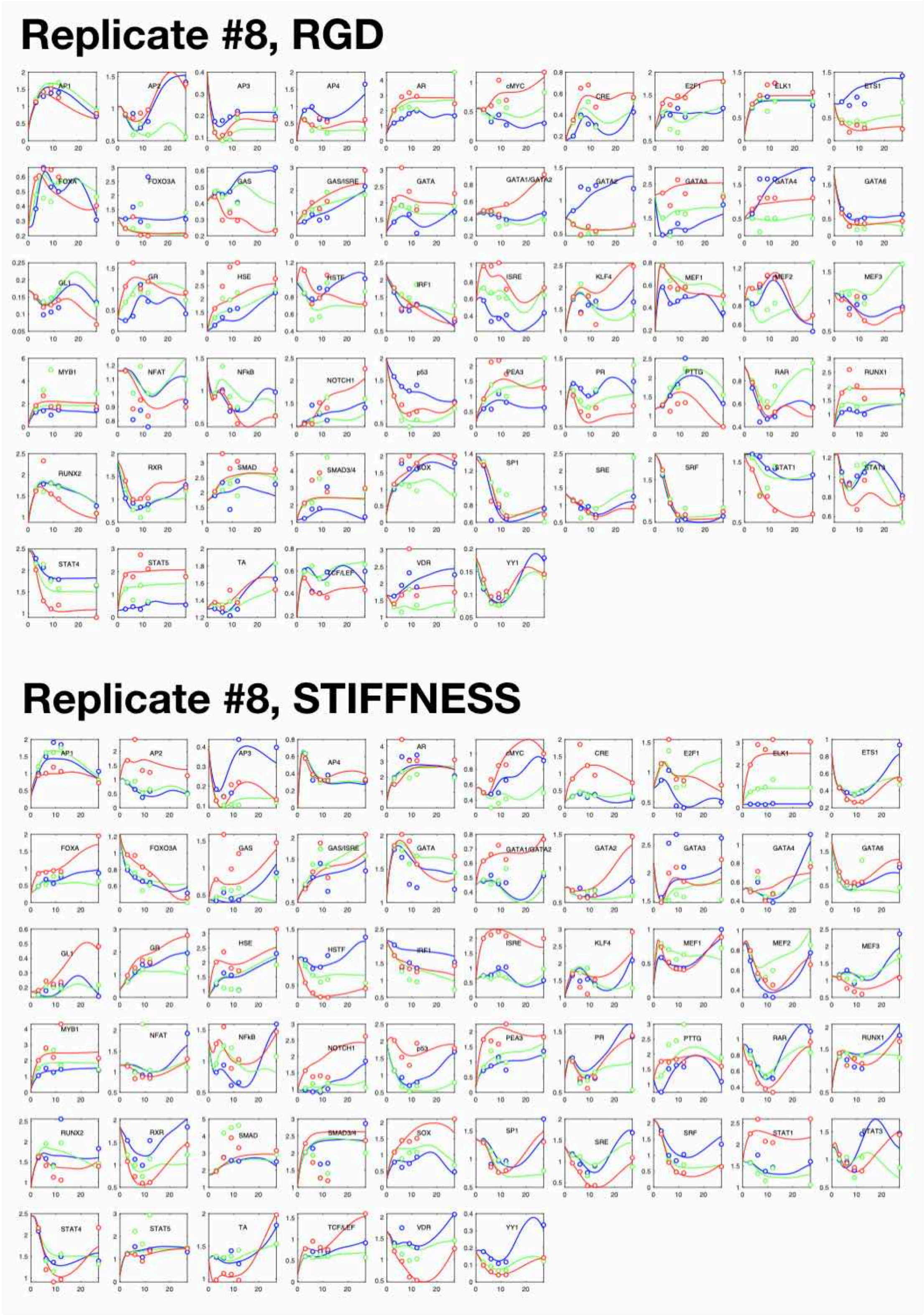
